# Nuclear actin regulates inducible transcription by enhancing RNA polymerase II clustering

**DOI:** 10.1101/655043

**Authors:** Mian Wei, Xiaoying Fan, Miao Ding, Ruifeng Li, Shipeng Shao, Yingping Hou, Shaoshuai Meng, Fuchou Tang, Cheng Li, Yujie Sun

## Abstract

Gene expression in response to external stimuli underlies a variety of fundamental cellular processes. However, how the transcription machinery is regulated under these scenarios is largely unknown. Here, we discover a novel role of nuclear actin in inducible transcriptional regulation using next-generation transcriptome sequencing and super-resolution microscopy. The RNA-seq data reveal that nuclear actin is required for the establishment of the serum-induced transcriptional program. Using super-resolution imaging, we found a remarkable enhancement of RNA polymerase II (Pol II) clustering upon serum stimulation and this enhancement requires the presence of nuclear actin. To study the molecular mechanisms, we firstly observed that Pol II clusters co-localized with the serum-response genes and nuclear actin polymerized in adjacent to Pol II clusters upon serum stimulation. Furthermore, N-WASP and Arp2/3 are reported to interact with Pol II, and we demonstrated N-WASP is required for serum-enhanced Pol II clustering. Importantly, using an optogenetic tool, we revealed that N-WASP phase-separated with the carboxy-terminal domain of Pol II and nuclear actin. In addition to serum stimulation, we found nuclear actin also essential in enhancing Pol II clustering upon interferon-γ treatment. Taken together, our work unveils nuclear actin promotes the formation of transcription factory on inducible genes, acting as a general mechanism underlying the rapid response to environmental cues.

---

Response to stimuli is a hallmark of life. Genetic regulation in response to stimuli determines the fundamental decision a living cell makes on proliferation, differentiation, motility, and cell death. At tissue and organism levels, precise control of genetic response has substantial impacts on organ development, immune responses, and neuronal plasticity, and may contribute to tumor progression upon its dysregulation^1-5^. Surprisingly, little is known about how the transcription machinery is regulated in these inducible transcription processes.

Nuclear actin is implicated in gene regulation through various mechanisms. Nuclear actin is present in chromatin remodeling complexes^6^; it interacts with all three classes of RNA polymerases for their full transcriptional activity^7-9^; it modulates the localization and activity of megakaryocytic acute leukemia (MAL), a coactivator of serum response factor (SRF)^10^; it associates with nascent transcripts through specific heterogeneous nuclear ribonucleoproteins (hnRNPs)^11, 12^; and it is involved in long-range chromatin organization^13-15^. Despite recent progress, the mechanisms whereby nuclear actin regulates gene expression remain unclear.

Recent studies have revealed that serum stimulation induces dynamic assembly of nuclear actin filaments, which regulates the SRF-MAL signaling pathway, a key pathway involved in inducible gene expression^16^. Noteworthily, we and others reported that serum stimulation also dramatically changes the dynamics of “transcription factories”^17, 18^, which are discrete RNA polymerase II (Pol II)-clustered foci implicated in gene regulation^19-23^. Thus, it would be intriguing to determine whether these two serum stimulation-dependent phenomena are physiologically connected. Here, we formulate a hypothesis that nuclear actin regulates inducible transcription by modulating the formation of transcription factories.

We test this hypothesis with a combination of next-generation transcriptome sequencing and super-resolution microscopy. We find that nuclear actin contributes to the establishment of the serum-induced transcriptional program by scaffolding larger, more active, and more long-lasting Pol II clusters upon serum stimulation. Similar observation is also obtained upon interferon-γ (IFN-γ) treatment. We demonstrate that nuclear actin promotes Pol II cluster formation through *de novo* nuclear actin polymerization induced by N-WASP/Arp2/3. Using an optoDroplet system, we observe N-WASP phase-separates with Pol II and nuclear actin. Taken together, our work unveils a novel role of nuclear actin in transcriptional modulation of inducible genes by promoting enhanced-level Pol II clustering, acting as a general mechanism underlying the rapid response to environmental cues.

## RESULTS

### Nuclear actin is required for the establishment of the serum-induced transcriptional program

To investigate the role of nuclear actin in inducible transcription activity, we interrogated whether the disruption of nuclear actin results in any change in the transcriptome upon serum stimulation. Commonly used pharmaceutical agents targeting actin, such as cytochalasin D and latrunculin B, lack the specificity to disrupt actin polymerization only in the nucleus. To overcome this difficulty and achieve high specificity towards nuclear actin, we fused actin mutant G13R, a dominant negative mutant that cannot polymerize^24^, with a nuclear localization signal (NLS) sequence and overexpressed the recombinant protein NLS-actin G13R (G13R) to specifically perturb the polymerization state of nuclear actin. We performed next-generation transcriptome sequencing on cells overexpressing G13R and those expressing GFP as the control, under two conditions: normal growth and serum stimulation (Supplementary Fig. 1). Serum stimulation modulates the transcription of specific genes called serum-response genes^1, 2^. In order to identify differentially expressed genes in response to serum, we defined a parameter termed “fold change” as the ratio of the transcription level under serum-stimulation condition to that under normal-growth condition. We found that in general, these serum-response genes exhibited a lower fold change in cells overexpressing G13R than in control cells overexpressing GFP (Fig. 1a). Furthermore, most of the serum-response genes in GFP-control cells were not present in cells overexpressing G13R (Fig. 1b). In contrast, for basal-level transcription under normal-growth condition, cells overexpressing G13R had a highly similar transcriptional profile to those expressing GFP, with only 38 differentially expressed genes (data not shown). To confirm the RNA-seq results, we selected nine canonical serum-response genes and two housekeeping genes *RPLP0* and *GAPDH* for qRT-PCR (*GAPDH* as the internal control). The qRT-PCR results were highly consistent with the RNA-seq data (Fig. 1c). Noteworthily, most canonical serum-response genes had a lower fold change in G13R-overexpressing cells than in GFP-control cells (Fig. 1c, left). Next, we sorted the up-regulated genes by the level of transcriptional change caused by G13R overexpression (Methods; Supplementary Table 1) and conducted functional annotation analysis of the top ∼30% ranked genes (Fig. 1d). These results indicate that nuclear actin is required for the serum-induced expression of specific transcriptional regulators. Taken together, these results demonstrate that nuclear actin is required for the establishment of the serum-induced transcriptional profile.

**Fig. 1.**
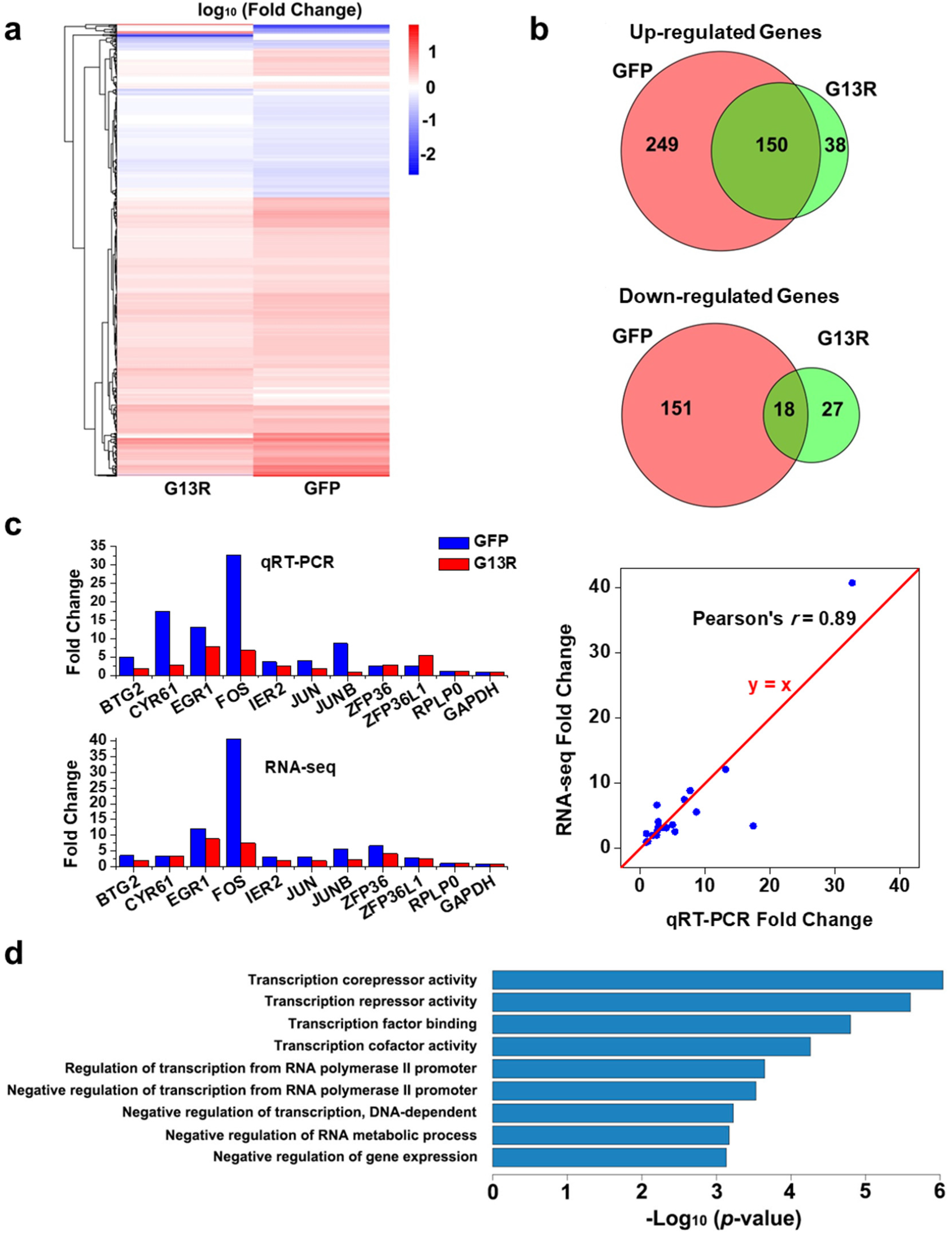
Nuclear actin is required for the establishment of the serum-induced transcriptional program. **a**, Heatmap showing the fold change for differentially expressed genes upon serum stimulation in cells overexpressing G13R and cells overexpressing GFP. **b**, Venn diagrams showing the numbers of differentially expressed genes. **c**, qRT-PCR conformation for nine canonical serum-response genes and two housekeeping genes. **d**, The functional annotations of the top ranked genes after sorting by the level of transcriptional change caused by actin mutant overexpression.

### Serum-response genes are localized within Pol II clusters upon serum stimulation for transcription

To probe the underlying mechanism for nuclear actin-dependent inducible transcription, we first investigate the transcriptional state of serum-responsive genes by imaging their localization relative to Pol II molecules *via* ImmunoFISH. Specifically, we found that Pol II molecules form clusters on two canonical serum-response genes—*FOS* and *JUNB*^1^, upon serum stimulation (Supplementary Fig. 2). On the contrary, for a control gene *PLA2G2E*, the percentage of alleles that colocalized with Pol II clusters remained virtually the same under both conditions (Supplementary Fig. 2). These results suggest that upon serum stimulation, inducible transcription of serum-response genes takes place in clusters of Pol II molecules.

In order to confirm that Pol II clusters are active transcription hubs, we investigated the spatial relationship between Pol II clusters and nascent RNA using two-color super-resolution imaging^25, 26^. To visualize Pol II, we constructed a human osteosarcoma cell line (U2OS) that stably expresses the Dendra2-fused catalytic subunit of Pol II (RPB1), replacing endogenous RPB1 as previously reported^17, 18^. We labeled nascent RNA by pulse incorporation of 5-ethynyl uridine (EU) and Click Chemistry-based conjugation of dyes to EU (Methods). We quantified the colocalization between Pol II clusters and nascent RNA by pair cross-correlation analysis^27^ and found that the colocalization was evidently higher than that of pixel-permutated images (Supplementary Fig. 3). Taken together, we reason that Pol II clusters form on specific serum-response genes for active transcription upon serum stimulation.

### Serum stimulation enhances the spatiotemporal dynamics of Pol II clustering

We previously reported that Pol II form clusters in the nucleus and that these clusters appeared and disappeared in an asynchronous manner^18^. Here, using live-cell epifluorescence microscopy, we counted the number of clusters under both normal-growth and serum-stimulation conditions. We found that cells had significantly more clusters under serum-stimulation condition than under normal-growth condition (Supplementary Fig. 4), as we previously reported^18^. These results show that serum stimulation enhances the formation of clusters that are large enough to be discerned by epifluorescence microscopy.

To quantify the spatiotemporal dynamics of Pol II clusters at higher resolution, we recorded live-cell photoactivated localization microscopy (PALM)^25^ images of this stable cell line (Fig. 2a). Pair correlation analysis^27^ confirmed Pol II clustering, with a correction for the photophysical properties of Dendra2 (Supplementary Fig. 5). We then used SR-Tesseler, a segmentation-based cluster analysis method^28^, to identify clusters and extract cluster characteristics (Fig. 2b,c). For the temporal dynamics of Pol II clusters, we applied the time correlated PALM (tcPALM) analysis as previously described^17^. In tcPALM, assembly of a Pol II cluster is represented by an abrupt increase of single-molecule localizations detected in a cluster, termed a “burst” (Methods; Supplementary Fig. 6). Using these analyses, we quantitatively compared Pol II clustering under three conditions: serum stimulation, serum deprivation, and normal growth. We found that cluster density, cluster area, and the number of localizations per cluster were indeed higher in serum-stimulated cells than under normal-growth condition (Fig. 2d, Supplementary Fig. 7a), in accord with our previous results^18^. Similarly, the number of localizations in a burst (burst size), the duration of a burst (burst lifetime), and the number of bursts per cluster obtained by tcPALM analysis were also larger in serum-stimulated cells (Fig. 2e, Supplementary Fig. 7b), consistent with a previous report^17^. In contrast, all of these cluster parameters in cells under serum-deprivation condition did not show significant difference from those under normal-growth condition (Fig. 2d,e, Supplementary Fig. 7), indicating that the serum-induced changes in Pol II clustering were due to serum stimulation rather than serum deprivation that preceded stimulation (Methods). These results suggest that more Pol II clusters form *de novo* upon serum stimulation, which accords with the on-demand model of transcription factory formation^17, 18, 22^, and that Pol II clusters become more active, grow larger, and contain more Pol II molecules upon serum stimulation.

**Fig. 2.**
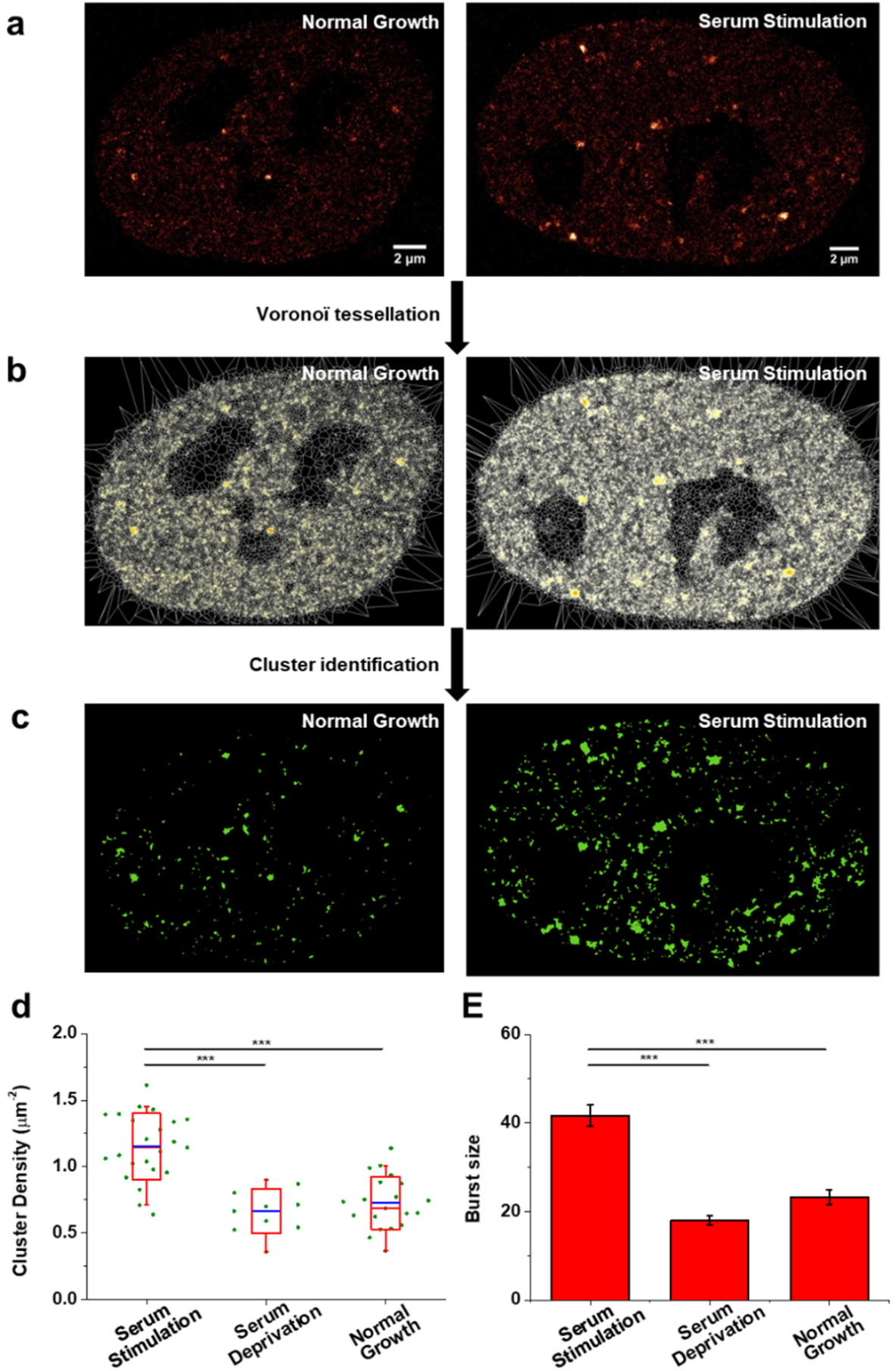
Enhanced Pol II clustering upon serum stimulation. **a**, Representative live-cell PALM images of Pol II under normal-growth condition (n = 35 cells) and under serum-stimulation condition (n = 39 cells). **b**, Voronoï tessellation of the images in (A) by SR-Tesseler. **c**, Images of Pol II clusters identified by SR-Tesseler. **d**, Cluster density under serum-stimulation (n = 23 cells), serum-deprivation (n = 10 cells), and normal-growth conditions (n = 20 cells). Each green dot is the mean for all Pol II clusters from a single nucleus. The blue line is the mean for the whole population of nuclei under the indicated condition. The red box shows the SD around the mean. The red line within the red box is the median. The whiskers show 5% and 95%. **e**, Burst size under serum-stimulation (n = 829 bursts of 104 clusters from 23 cells), serum-deprivation (n = 174 bursts of 61 clusters from 10 cells), and normal-growth conditions (n = 306 bursts of 111 clusters from 20 cells). Data are shown as mean ± SEM. Statistical significance was determined by one-way ANOVA with Tukey–Kramer test. \**p*< 0.05, \*\**p*< 0.01, and \*\*\**p*< 0.001.

### Nuclear actin is required for serum-enhanced Pol II clustering

As serum stimulation drastically enhanced Pol II clustering dynamics, we investigated whether nuclear actin was required for this enhanced-level Pol II clustering. To address this question, we overexpressed NLS-actin mutants (G13R, R62D, and S14C)^24^ to perturb the polymerization state of nuclear actin. G13R and R62D are two dominant negative mutants that cannot polymerize, while S14C is a mutant that is more likely to polymerize than wild-type actin. Indeed, overexpression of S14C promoted a massive assembly of nuclear actin filaments, while G13R and R62D did not have such effects (Supplementary Fig. 8). With overexpression of nuclear actin mutants, serum stimulation was no longer able to promote the formation of Pol II clusters, which reduced to a level similar to that in cells under normal-growth condition (Fig. 3, Supplementary Fig. 9). Note that without serum stimulation, there was no significant difference in any of the cluster parameters in cells overexpressing these actin mutants compared to those under normal-growth condition (Supplementary Fig. 10). These results suggest that dynamic polymerization and depolymerization of nuclear actin is required for enhanced-level Pol II clustering upon serum stimulation, but is dispensible for basal-level Pol II clustering under normal-growth condition. In addition to actin mutants, we also carried out spatial knockdown of actin in the nucleus by overexpressing exportin 6 (XPO6) and studied its effects on Pol II clustering. XPO6 specifically exports actin-profilin complexes into the cytoplasm^29^. As expected, the fluorescent levels of nuclear actin were dramatically lower in XPO6-overexpressing cells than in cells that did not overexpress XPO6 (Supplementary Fig. 11). Upon depletion of nuclear actin by overexpressing XPO6, we found that serum stimulation was no longer able to promote the formation of Pol II clusters, which reduced to a level similar to that in cells under normal-growth condition (Fig. 4, Supplementary Fig. 12). Note that without serum stimulation, there was no significant difference in any of the cluster parameters in XPO6-overexpressing cells compared to the control cells under normal-growth condition (Fig. 4). These results confirm that nuclear actin is required for enhanced-level Pol II clustering, but dispensable for basal-level Pol II clustering.

**Fig. 3.**
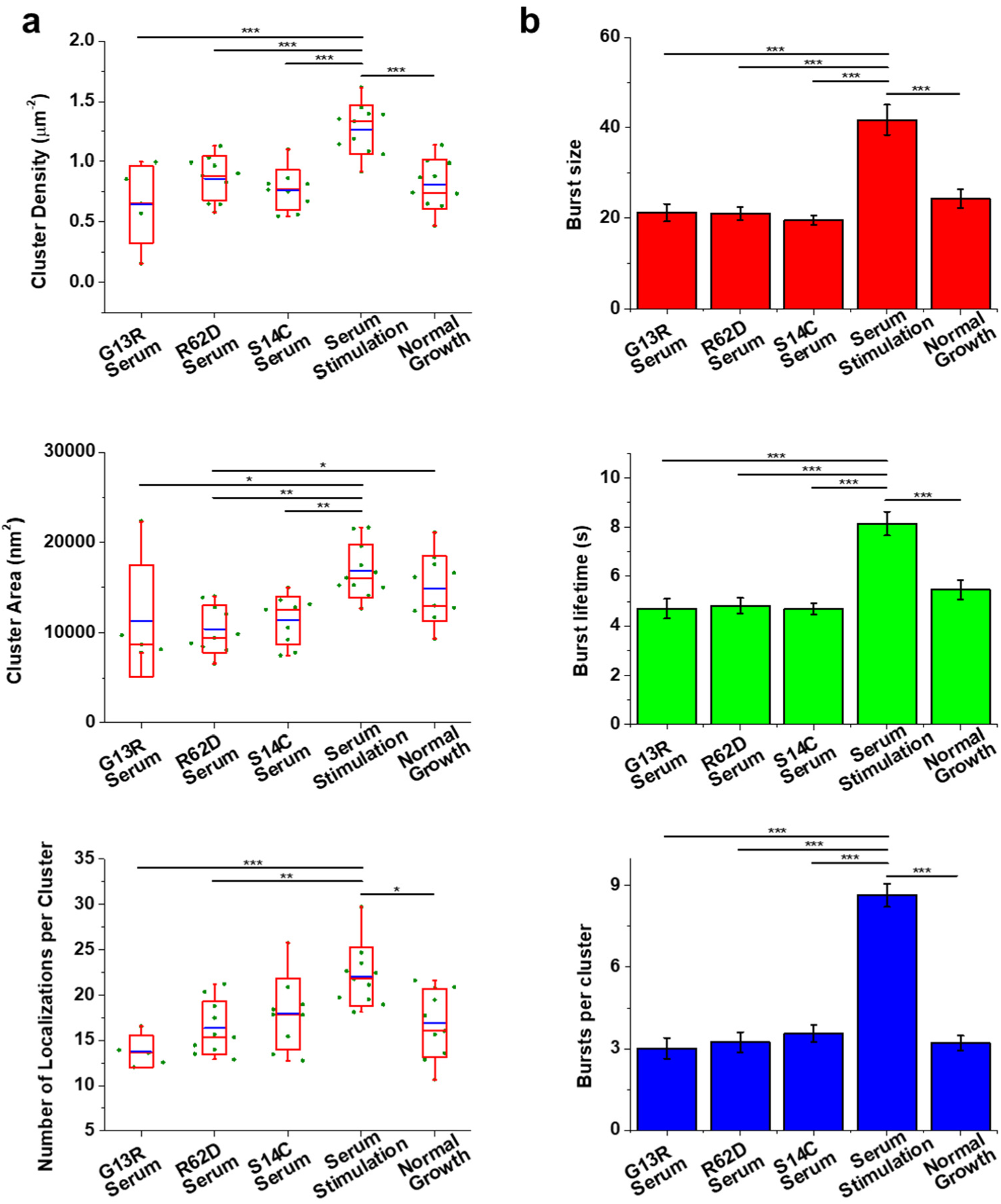
Overexpression of actin mutants abolishes serum-enhanced Pol II clustering. **a**, Cluster density, cluster area, and the number of localizations per cluster in cells overexpressing different actin mutants under serum-stimulation condition (n = 5, 10, and 9 cells, respectively, from left to right), cells under serum-stimulation condition (n = 11 cells), and cells under normal-growth condition (n = 10 cells). Each green dot is the mean for all Pol II clusters from a single nucleus. The blue line is the mean for the whole population of nuclei under the indicated condition. The red box shows the SD around the mean. The red line within the red box is the median. The whiskers show 5% and 95%. **b**, Burst size, burst lifetime, and the number of bursts per cluster in cells overexpressing different actin mutants under serum-stimulation condition (n = 163 bursts of 54 clusters from 5 cells, 230 bursts of 71 clusters from 10 cells, and 271 bursts of 76 clusters from 9 cells, respectively, from left to right), cells under serum-stimulation condition (n = 432 bursts of 50 clusters from 11 cells), and cells under normal-growth condition (n = 192 bursts of 60 clusters from 10 cells). Data are shown as mean ± SEM. Statistical significance was determined by one-way ANOVA with Tukey–Kramer test. \**p*< 0.05, \*\**p*< 0.01, and \*\*\**p*< 0.001.

To eliminate the possibility that the transfection procedure interfered with the enhanced-level Pol II clustering upon serum stimulation, we overexpressed GFP as the control to investigate its effect on Pol II clustering. We found that GFP overexpression hardly affected either the spatial organization or the temporal dynamics of Pol II clusters under both serum-stimulation and normal-growth conditions (Supplementary Fig. 13). Overexpression of NLS-wild-type actin did not affect the spatial organization but interfered with the temporal dynamics of Pol II clusters under serum-stimulation condition (Supplementary Fig. 14). This complex phenotype is possibly due to the fact that the amount of nuclear actin is inherently coupled to the polymerization state of nuclear actin. Aside from increasing the amount of nuclear actin, overexpression of NLS-actin inevitably changed the ratio of F-actin to G-actin and thus perturbed actin dynamics in the nucleus, as shown in a recent study reporting that overexpression of EGFP-NLS-actin formed large nuclear actin filaments that could be stained by phalloidin^30^.

### Nuclear actin filaments form in adjacent to Pol II clusters upon serum stimulation

To investigate how nuclear actin regulates Pol II clustering upon serum stimulation, we studied the spatial relationship between nuclear actin and Pol II clusters using two-color super-resolution imaging. Imaging nuclear actin has long been a challenge because the signal of cytoplasmic actin overwhelms that of nuclear actin due to its relatively low abundance^31^. Moreover, the currently available live-cell imaging probes for nuclear actin interfere with actin dynamics and result in artifacts^32-34^. To overcome these challenges, we used an extraction procedure to remove most of the cytoplasmic substances and then labeled actin by dye-conjugated phalloidin, a well-established F-actin probe, in fixed cells (Methods). Nuclear actin appeared as dense clusters in the nucleoplasm^32, 35^ and was also enriched in nucleoli and nuclear membrane (Fig. 5a). The enrichment of nuclear actin in nucleoli is consistent with its role in Pol I transcription^9, 36, 37^. Nuclear membrane-associated cytoplasmic actin is tethered to the nuclear lamina *via* the linker of nucleoskeleton and cytoskeleton complex^38^. Regarding the spatial relationship between nuclear actin and Pol II clusters, both visual inspection (Fig. 5a) and quantitative analysis (Fig. 5b) indicate that nuclear actin clusters evidently colocalized with Pol II clusters at a much higher level under serum-stimulation condition than that under normal-growth condition. In addition, this serum-induced colocalization was significantly higher than the random level (Supplementary Fig. 15). These results suggest that nuclear actin filaments scaffold Pol II clusters through protein-protein interactions upon serum stimulation.

**Fig. 4.**
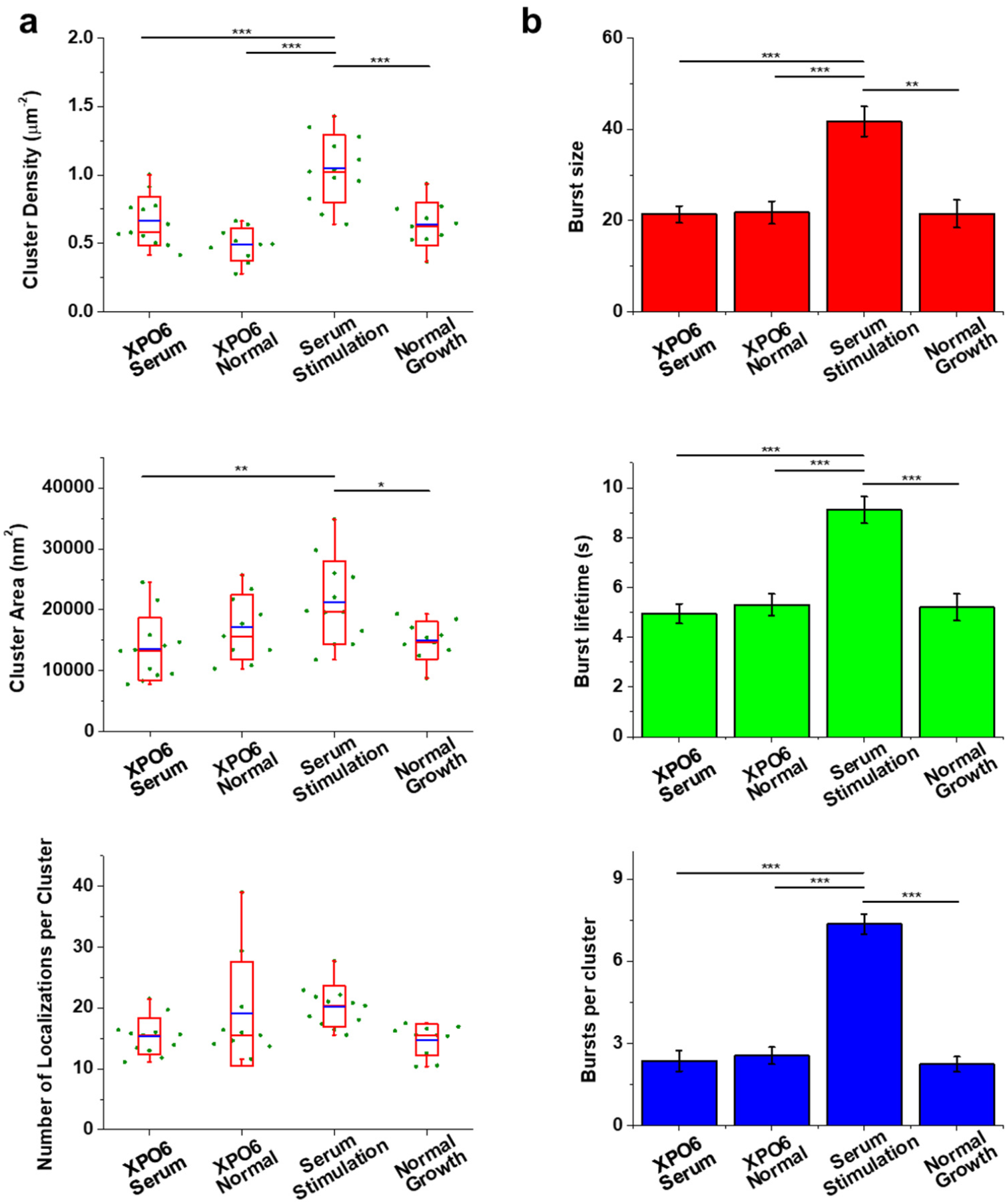
Overexpression of XPO6 abolishes serum-enhanced Pol II clustering. **a**, Cluster density, cluster area, and the number of localizations per cluster in cells overexpressing XPO6 under serum-stimulation (n = 12 cells) and normal-growth (n = 10 cells) conditions, cells under serum-stimulation condition (n = 12 cells), and cells under normal-growth condition (n = 10 cells). Each green dot is the mean for all Pol II clusters from a single nucleus. The blue line is the mean for the whole population of nuclei under the indicated condition. The red box shows the SD around the mean. The red line within the red box is the median. The whiskers show 5% and 95%. **b**, Burst size, burst lifetime, and the number of bursts per cluster in cells overexpressing XPO6 under serum-stimulation (n = 144 bursts of 61 clusters from 12 cells) and normal-growth (n = 125 bursts of 49 clusters from 10 cells) conditions, cells under serum-stimulation condition (n = 397 bursts of 54 clusters from 12 cells), and cells under normal-growth condition (n = 114 bursts of 51 clusters from 10 cells). Data are shown as mean ± SEM. Statistical significance was determined by one-way ANOVA with Tukey–Kramer test. \**p*< 0.05, \*\**p*< 0.01, and \*\*\**p*< 0.001.

**Fig. 5.**
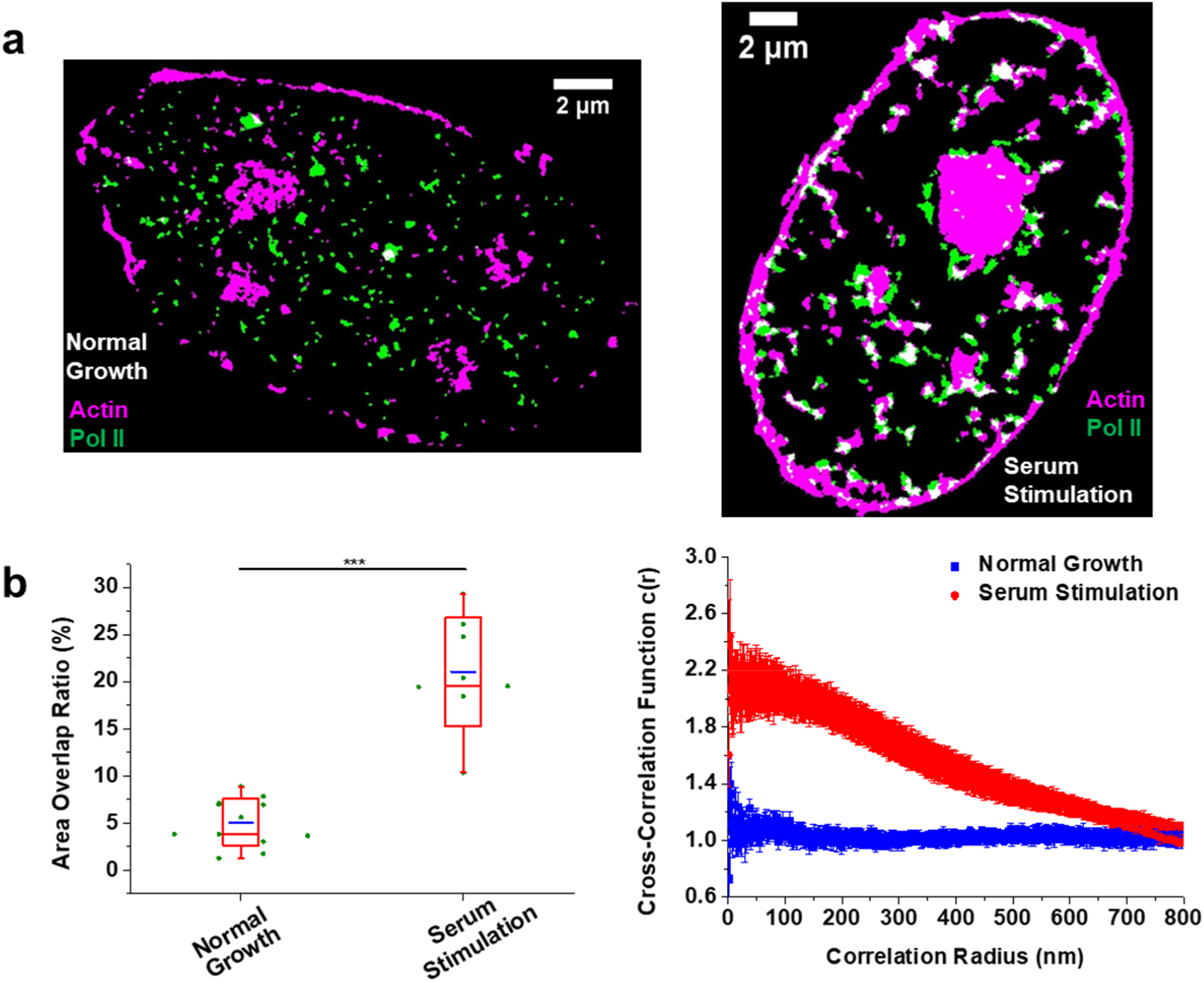
Nuclear actin filaments colocalize with Pol II clusters upon serum stimulation. **a**, Representative cluster images obtained by SR-Tesseler showing actin (magenta) and Pol II (green) in cells under normal-growth (n = 12 cells) and serum-stimulation (n = 8 cells) conditions. **b**, Quantification of colocalization by calculating the area overlap ratio (left) and analyzing the pair cross-correlation (right). Left: Each green dot is the area overlap ratio fora single nucleus. The blue line is the mean for the whole population of nuclei under the indicated condition. The red box shows the SD around the mean. The red line within the red box is the median. The whiskers show 5% and 95%. Statistical significance was determined by two-tailed *t*-test. \**p*< 0.05, \*\**p*< 0.01, and \*\*\**p*< 0.001. Right: Data are shown as mean ± SEM. (n = 12 cells under normal-growth condition, n = 8 cells under serum-stimulation condition)

### N-WASP-induced nuclear actin polymerization is required for serum-enhanced Pol II clustering

Previous studies have found that both the actin nucleator Arp2/3 and its activating protein N-WASP localized in the nucleus and play a crucial role in Pol II-dependent transcription^39, 40^. Importantly, N-WASP is found to physically associate with Pol II through its interaction with NonO, which binds the carboxy-terminal domain (CTD) of Pol II. It is also demonstrated that the regulation of Pol II-dependent transcription by N-WASP depends on *de novo* formation of actin filaments induced by N-WASP, rather than the pre-existing actin filaments^39^. A following work confirms that N-WASP-induced nuclear actin polymerization is mediated by the Arp2/3 complex, which also physically associates with Pol II^40^. To investigate whether the nuclear actin filaments induced by N-WASP and Arp2/3 are responsible for serum-enhanced Pol II clustering, we constructed a nuclear-localized dominant negative mutant of N-WASP (NLS-CA)^39^ and studied the effect of N-WASP on the temporal dynamics of Pol II clusters. We found that upon overexpression of NLS-CA, serum stimulation was no longer able to promote the formation of Pol II clusters. The burst size, burst lifetime, and the number of bursts per cluster all reduced to a level similar to that in cells under normal-growth condition (Fig. 6), and also similar to that in cells overexpressing actin mutants (Fig. 3b) or XPO6 (Fig. 4b). These results indicate that *de novo* nuclear actin polymerization through the N-WASP-Arp2/3 pathway is required for enhanced Pol II clustering upon serum stimulation.

**Fig. 6.**
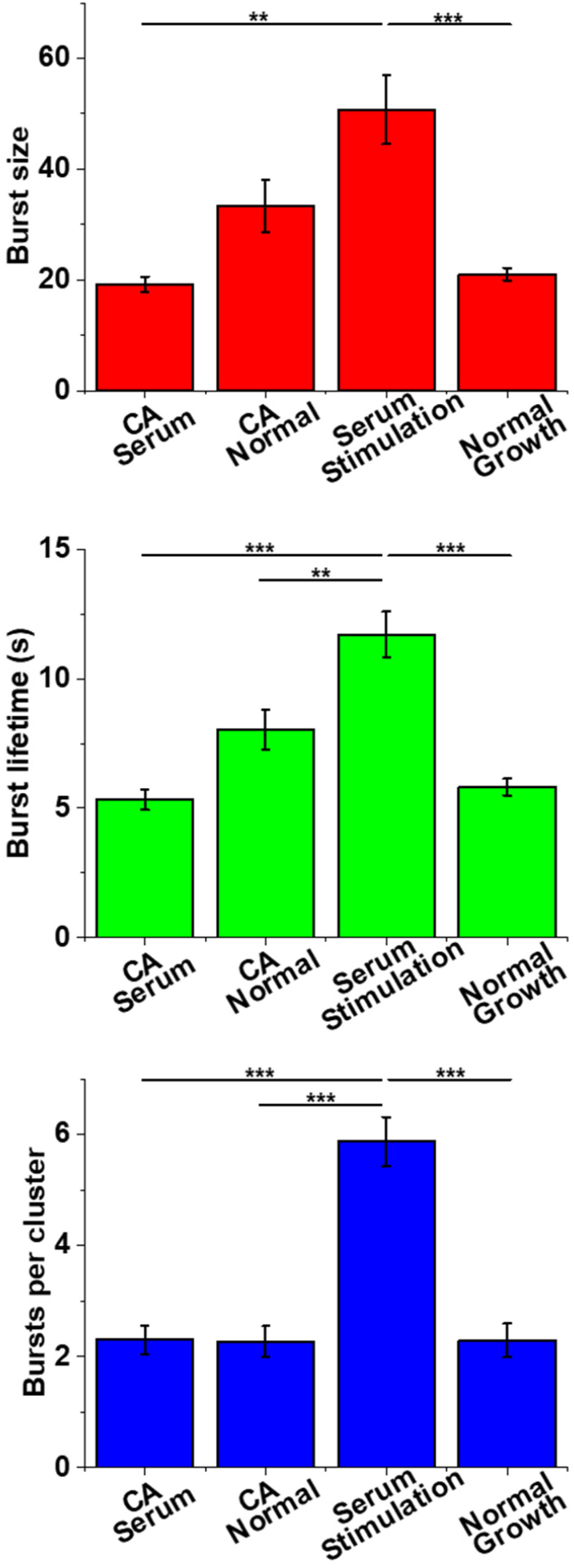
Overexpression of a dominant negative mutant of N-WASP (NLS-CA) interferes with the temporal dynamics of Pol II clusters. Burst size, burst lifetime, and the number of bursts per cluster in cells overexpressing NLS-CA under serum-stimulation (n = 115 bursts of 50 clusters from 10 cells) and normal-growth (n = 152 bursts of 67 clusters from 13 cells) conditions, cells under serum-stimulation condition (n = 323 bursts of 55 clusters from 11 cells), and cells under normal-growth condition (n = 149 bursts of 65 clusters from 13 cells). Data are shown as mean ± SEM. Statistical significance was determined by one-way ANOVA with Tukey–Kramer test. \**p* < 0.05, \*\**p* < 0.01, and \*\*\**p* < 0.001.

As both N-WASP and Pol II contain low-complexity domains for liquid-liquid phase separation^41-44^, we hypothesized that N-WASP-induced nuclear actin scaffolds Pol II clusters through a phase separation mechanism. To test this hypothesis, we constructed an optoDroplet system^45^ by fusing N-WASP to Cry2 for light-activated control of intracellular phase separation. Upon light activation, N-WASP phase-separated with the CTD of RPB1 and nuclear actin into liquid droplets in the nucleus (Supplementary Fig. 16). These results suggest a phase separation mechanism by which N-WASP-induced nuclear actin scaffolds Pol II clusters.

### Nuclear actin is also required for enhanced-level Pol II clustering upon IFN-γ treatment

To test if nuclear actin-enhanced Pol II clustering acts as a general mechanism for cells in response to environmental cues, we investigated the effect of nuclear actin in another inducible transcription system. IFN-γ is a crucial cytokine for innate and adaptive immunity^46^. Treatment of interferons was previously found to cause the recruitment of nuclear actin to the promoter regions of interferon-inducible genes^8^. For IFN-γ induced gene expression, we found that nuclear actin indeed promoted enhanced-level Pol II clustering upon IFN-γ treatment in a similar manner to serum stimulation (Supplementary Fig. 17), suggesting the general role of nuclear actin in regulating Pol II clustering and gene expression in response to external stimuli.

## DISCUSSION

Both nuclear actin and transcription factories have been under fierce debate^31, 47^. In this study, we propose a mechanism in which nuclear actin regulates inducible transcription by promoting enhanced-level Pol II clustering in response to stimuli. Specifically, we investigated the effects of nuclear actin on Pol II clustering upon serum stimulation as well as IFN-γ treatment. Under normal-growth condition, single Pol II molecules assemble into clusters for basal-level transcription, which is independent of nuclear actin dynamics (Supplementary Fig. 4). In contrast, under stimulation condition, Pol II molecules assemble into larger, more dynamic, and more long-lasting clusters which requires dynamic polymerization and depolymerization of nuclear actin (Fig. 3, Fig. 7a). This work connects nuclear actin dynamics with Pol II transcription factories by unveiling the mechanism underlying rapid modulation of gene expression in response to environmental cues, adding insights on actin-related signaling pathways (Fig. 7b).

**Fig. 7.**
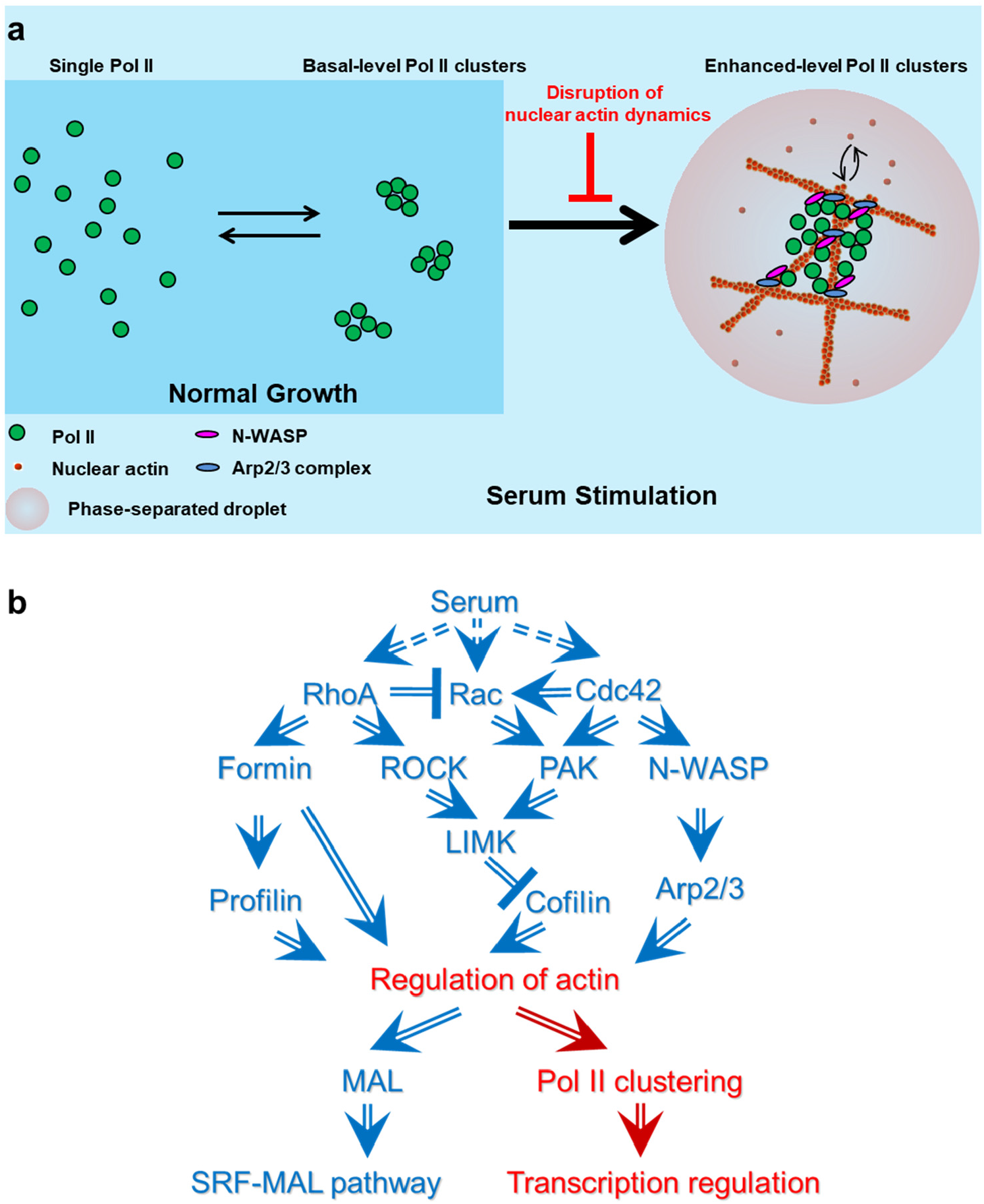
Model that Nuclear Actin Dynamics Regulate Pol II Clustering in Response to Serum. **a**, Pol II molecules assemble into basal-level clusters for transcription under normal-growth condition, which is independent of nuclear actin dynamics. Upon serum stimulation, Pol II molecules aggregate into enhanced-level clusters for serum-induced transcription, which requires dynamic polymerization and depolymerization of nuclear actin. **b**, Actin-related signaling pathway diagram showing what is known (blue) and what is proposed and investigated in this study (red arrows).

The regulation of Pol II clustering by nuclear actin may promote the formation of the preinitiation complex and transcription initiation in response to stimuli. This conclusion is based on previous studies by others and us reporting that inhibitors of transcription elongation do not reduce the dynamics or density of Pol II clusters^17, 18^, suggesting Pol II clustering takes place in the transcription initiation stage. Thus, our model is compatible and complementary to previous models that monomeric actin contributes to chromatin remodeling by mediating the binding of the chromatin remodeling complex INO80 to chromatin^48^ and nuclear actin facilitates transcription elongation by assembling at the Ser2-phosphorylated Pol II CTD with hnRNP U to recruit the histone acetyltransferase PCAF^7, 12, 49^. Besides, a recent study investigated the role of nuclear actin in basal-level transcription and Pol II clustering^30^, complementing our work in inducible transcription under external stimuli.

Our data suggest that nuclear actin may work as a dynamic scaffold to foster the formation of enhanced-level Pol II clusters through a phase separation mechanism (Fig. 7a, Supplementary Fig. 16). The phase separation of N-WASP through its proline-rich motif has been shown to facilitate Arp2/3-branched actin polymerization^43^. The phase separation of Pol II through the CTD of the RPB1 subunit is regulated by CTD phosphorylation and plays a role in promoter escape and transcription elongation^41, 42, 44^. Importantly, N-WASP and Arp2/3 physically associates with Pol II and this association is involved in Pol II transcription^39, 40^. Using an optoDroplet system, we demonstrated that N-WASP phase-separated with the CTD of Pol II and nuclear actin upon light activation, mimicking the inducible formation of enhanced-level Pol II clusters upon serum stimulation (Supplementary Fig. 16). Our model that enhanced-level Pol II clusters phase-separate upon serum stimulation (Fig. 7a) is consistent with a recent report showing transient interactions between transcription factors and Pol II without detectable phase separation under normal-growth condition^50^.

The precise molecular mechanisms of how nuclear actin scaffolds Pol II clusters on specific genes in response to stimuli still remain to be elucidated. Firstly, it is intriguing to understand how nuclear actin scaffolding, co-phase separating with Pol II clusters can function as a general mechanism to promote transcription factory formation in response to different stimuli, in this work serum and IFN-γ, respectively. Several studies have suggested a transcription factor-dependent transcription factory formation model in which genes regulated by the same transcription factor and cofactors tend to cluster in the same transcription factory^21, 51^. We speculate that following the transcription factor-dependent clustering of genes and Pol II molecules, Pol II can recruit N-WASP and Arp2/3, which subsequently activate nuclear actin polymerization and enhance Pol II clustering in a positive feedback manner through multi-valency interaction and phase separation^43^. Detailed investigation of this speculation and the roles of other factors such as NonO and PSF^39, 40^ and phosphorylation of the CTD of Pol II^41, 42, 44^ would be of great importance for understanding the precise mechanisms. Secondly, technological advancements, especially in imaging, would greatly benefit this investigation^52, 53^. Recent studies, together with our unpublished data, indicate that various live-cell imaging probes for nuclear actin induce artifacts^32-34^. The development of methods that allow artifact-free, high-contrast, and live-cell imaging of nuclear actin would aid in dissecting the details of gene regulation by nuclear actin.

## METHODS

Methods, including statements of data availability and any associated accession codes and references, are available in the online version of this paper.

*Note: Supplementary Information is available in the online version of the paper*.

## ACKNOWLEDGMENTS

We thank X. Zhuang (Harvard University), C. Wu (Peking University), K. Xu (University of California, Berkeley) for discussion, H. Babcock (Harvard University) for help with optical setup of the STORM microscopy, B. Huang (University of California, San Francisco) for Insight3 software and discussion, I. I. Cisse (Massachusetts Institute of Technology) for help with tcPALM experiments, F. Levet (University of Bordeaux) for help with SR-Tesseler analysis, W. Du (Tsinghua University) for help with cell biology experiments, N. Wang (University of California, Los Angeles) for help with bioinformatic analysis, W. Guo (University of Pennsylvania) for the gift of the GST-CA plasmid, and Z. Jiang (Peking University) for the gift of IFN-γ. We thank H. Lv, C. Shan, and the Core Facilities at School of Life Sciences, Peking University for imaging support. This work is supported by grants from the National Science Foundation of China 21573013, 21390412, 31271423, and 31327901, 863 Program SS2015AA020406 and CAS Interdisciplinary Innovation Team for Y.S.

## AUTHOR CONTRIBUTIONS

Y.S. and M.W. conceived the project and designed the experiments. M.W. performed the imaging experiments and data analysis with the help of M.D. X.F. performed RNA-seq and qRT-PCR experiments. R.L. conducted bioinformatic analysis. Y.H., M.W., and S.M. prepared the samples for ImmunoFISH. S.S. performed the optoDroplet assay. M.D. assisted with plasmid construction. M.W. and Y.S. wrote the manuscript, with inputs from X.F., R.L., Y.H., S.S., and M.D. All authors participated in discussion and editing of the manuscript.

## COMPETING FINANCIAL INTERESTS

The authors declare no competing financial interests.

## METHODS

### Cells and Plasmids

The original U2OS cell line was purchased from the Cell Bank of Chinese Academy of Sciences. A stable cell line was constructed to express Dendra2-RPB1 replacing endogenous RPB1 as previously described^17, 18^. The cell line was cultured in low-glucose DMEM (Life Technologies, 10567-014) supplemented with 10% FBS (Life Technologies, 10099-141) and 1% penicillin/streptomycin (Life Technologies, 15140-122). Cells were tested for the absence of mycoplasma using LookOut Mycoplasma PCR Detection Kit (Sigma-Aldrich, MP0035). No cell lines used in this study were listed in the database of commonly misidentified cells lines maintained by ICLAC and NCBI Biosample. We did not attempt to authenticate the cell lines. Transfections were conducted using FuGENE 6 (Promega, E2691) for imaging experiments or Amaxa Cell Line Nucleofector Kit V (Lonza Amaxa, VCA-1003) for RNA-seq experiments, according to the supplier’s protocols.

The plasmids co-expressing actin mutants and GFP were constructed as follows. Actin mutant fragments were PCR-amplified from wild-type β-actin with mutagenic primers. Thereafter, GFP, P2A, NLS, and actin mutant fragments were fused together by overlap PCR. Then the GFP-P2A-NLS-actin mutant fragments were inserted into the pcDNA 3.1(+) vector. The plasmid co-expressing wild-type actin and GFP was constructed in the same way. The plasmid coding for XPO6-mCerulean was constructed as follows: the XPO6 gene was PCR-amplified from human cDNA, fused with mCerulean by overlap PCR, and inserted into the pcDNA 3.1(+) vector. The plasmid expressing YFP-actin was constructed by inserting the PCR-amplified β-actin gene sequence into the pYFP-C1 vector. The plasmid co-expressing NLS-CA (a nuclear-localized dominant negative mutant of N-WASP) and GFP was constructed as follows: the CA sequence (the C-terminal portion of the N-WASP VCA domain) was PCR-amplified from the GST-CA plasmid (a gift from Wei Guo); GFP, P2A, NLS, and CA fragments were fused together by overlap PCR; and then the GFP-P2A-NLS-CA fragment was inserted into the pcDNA 3.1(+) vector.

### Serum Deprivation and Stimulation Experiments

Cells under serum deprivation condition were obtained by replacing the normal culture medium with serum-free DMEM containing 1% penicillin/streptomycin and culturing overnight. Serum-free L-15 medium (Life Technologies, 21083-027) was used for imaging under serum deprivation condition. Cells under serum-stimulation condition were cultured in the serum-free medium overnight and then stimulated with L-15 medium containing 20% FBS for 30 min before imaging. As a control, cells under normal-growth condition were grown in the normal culture medium and then transferred to L-15 medium containing 10% FBS right before imaging. Transfections were performed one day before culturing cells in the serum-free medium overnight.

### IFN-γ Treatment

Human IFN-γ (Sino Biological Inc., 11725-HNAS) was a gift from Z. Jiang lab (Peking University). Cells were left untreated or treated with IFN-γ (500 IU/ml) for 16 h before imaging. Transfections were performed one day before the treatment.

### Phase Separation Using an optoDroplet System

N-WASP was fused with Cry2-mCherry, which can form clusters containing several molecules upon blue light illumination. This process can be strengthened when Cry2-mCherry was fused with proteins containing intrinsically disorder domain (IDR). Cells were transfected with plasmids encoding Cry2-mCherry-N-WASP and EGFP-NLS-RPB1CTD. After 24 h, cells were illuminated by a LED light source for 20 minutes. Then cells were fixed using 4% PFA for 15 minutes. During the fixation process, the blue light was kept illuminating on cells to avoid the disassembly of phase-separated droplets. Cells were washed twice with pre-warmed phosphate-buffered saline, pH 7.4 (PBS). Then cells were permeabilized with 0.1% Triton X-100 in PBS for 15 minutes. Cells were stained in PBS solution containing 165 nM Alexa Fluor 647 Phalloidin (Molecular Probes, A22287) and 1% bovine serum albumin (BSA) for 30 minutes. Cells were stained with DAPI for 5 minutes and washed 3 times with PBS. Images were taken under a spinning disc confocal microscopy.

### Characterization of Actin Mutants

Cells cultured on glass-bottom dishes were left untreated or transfected with the plasmids encoding actin mutants. After 24 h, the cells were washed twice with PBS, fixed with 4% paraformaldehyde in PBS for 20 min, washed twice with PBS, and permeabilized with 0.5% Triton X-100 in PBS for 10 min. The cells were then washed twice with PBS and stained with 0.5 μM Alexa Fluor 647 Phalloidin in PBS for 30 min. The cells were washed twice with PBS and stained with 500 ng/ml DAPI for 2min. Thereafter, the cells were washed three times with PBS and immediately mounted for epifluorescence imaging.

### RNA-seq

After cells in a 6-well plate were rinsed with DPBS (Gibco, 14190-250), 1 ml of TRIzol LS Reagent (Ambion, 10296010) was added to one well. The cells were resuspended by pipetting and transferred into a 1.5-ml nuclease-free tube. The cell suspension was vigorously vortexed for 30 sec and then maintained at room temperature for 5 min. Thereafter, 200 μl of chloroform-isoamyl alcohol (24:1, Sigma-Aldrich, C0549-1PT) was added to the cell lysate and mixed by pipetting. The mixture was maintained at room temperature for 5 min and then centrifuged at 12,000 × *g* for 15 min at 4 °C. The top aqueous phase containing the total RNA was transferred to a new 1.5-ml tube. An equal volume of isopropanol (Beijing Chemical Works) was added, vortexed, and then maintained at room temperature for 10 min. The tube was centrifuged at 12,000 × *g* for 10 min at 4 °C to obtain the pellet, which was then washed with 1 ml of 80% ethanol (Sigma-Aldrich, 24102) diluted with DEPC-treated water (Ambion, AM9915G). After the pellet was dried, the total RNA was dissolved in 20 μl of RNase-free water. Approximately, 500 ng of total RNA (quantified by Nanodrop 2000) was used as the starting material for mRNA separation (NEBNext Poly(A) mRNA Magnetic Isolation Module, NEB, E7490L) and library preparation (NEBNext Ultra RNA Library Prep Kit for Illumina, NEB, E7530L). All steps were performed following the supplier’s protocols, except that the volume of each solution was half of the recommended volume for each reaction. The libraries were quality-tested for the appropriate fragment distribution and then sequenced on Illumina platform with 150 bp pair-end reads.

### Bioinformatic Analysis of RNA-seq

Whole transcriptome reads were aligned using the TopHat (v2.0.13)^54^ with the Ensembl Homo_sapiens_Ensembl_GRCh37.gtf as a reference (http://www.ensembl.org). In total, we sequenced 10 Mb and 13.7 Mb of 150 bp read pairs for duplicates of cells overexpressing actin mutant G13R under normal-growth condition, 10.9 Mb and 13.9 Mb of 150 bp read pairs for duplicates of cells overexpressing actin mutant G13R under serum-stimulation condition, 10.2 Mb and 16 Mb of 150 bp read pairs for duplicates of cells overexpressing GFP under normal-growth condition, and 14 Mb and 10 Mb of 150 bp read pairs for duplicates of cells overexpressing GFP under serum-stimulation condition. The transcriptional analysis was performed using Cufflinks (v2.2.1)^54^. The RPKM method (reads per kb of transcript per million mapped sequence reads)^55^ was used for normalizing gene counts. We calculated the fold change and *p*-value (*t*-test) of RPKM between the samples under serum-stimulation condition and the samples under normal-growth condition. Differentially expressed genes were screened using a fold change >2 and an absolute difference of FPKM > 5 between the samples under serum-stimulation condition and the samples under normal-growth condition (Supplementary Table 1). To determine the level of transcriptional change caused by actin mutant overexpression, we defined the fold change ratio of a gene as the ratio of its fold change in GFP-control cells to its fold change in cells overexpressing G13R. All the up-regulated genes in either GFP-control cells or cells overexpressing G13R (437 genes in total) were sorted by descending fold change ratio (Supplementary Table 1). The up-regulated genes were then divided into three groups: high fold-change-ratio group (140 genes), medium fold-change-ratio group (140 genes), and low fold-change-ratio group (157 genes). Functional annotation analysis was conducted for these three gene group using DAVID (https://david.ncifcrf.gov)^56^. The first annotation cluster of each group was presented in Supplementary Table 1. For the high fold-change-ratio group, the functional annotations in the first annotation cluster with a *p*-value below 0.001 were presented in Fig. 1D. The R density plot package (R Core Team. 2015. R: A language and environment for statistical computing. R Foundation for Statistical Computing, Vienna, Austria. https://www.R-project.org/) was used to generate the heatmaps, scatter plots, and Venn diagrams. Principal component analysis was conducted using the princomp function in the R graphics package.

### qRT-PCR

Isolation of total RNA was performed as the procedures in RNA-seq. Approximately, 200 ng of total RNA (quantified by Nanodrop 2000) was used for reverse transcription in a volume of 19 μl with oligo-d(T) primers using Superscript III reverse transcriptase (Invitrogen, 18080-044). After reverse transcription at 50 °C for 1 hour and 70 °C for 15 min to inactivate the enzyme, 1 μl of RNase H (Invitrogen, 18021-071) was added into each reaction tube to digest RNA. Thereafter, 180 μl of nuclease-free water was added to dilute the cDNA to 1 ng/μl. Of this diluted DNA, 1.5 μl was used for each 10 μl quantitative PCR reaction. The qPCR reaction was run on an ABI 7500 system using SYBR Green (Roche, 13396700) with the following protocol: 95 °C for 10 min, and 40 cycles at 95 °C for 5 sec and 60 °C for 30 sec. Besides the nine serum-response genes we selected from the RNA-seq data, we also selected two housekeeping genes, *GAPDH* and *RPLP0*, which served as controls. By normalizing the Ct values of each target gene to that of *GAPDH* for each sample, we calculated the fold change of a gene as dividing the ΔCt of the gene in cells under serum-stimulation condition by that under normal-growth condition.

### Probe Design and Synthesis for ImmunoFISH

Probe design and synthesis was conducted as previously described^57^. We designed human genomic libraries of PCR primer pairs delimiting amplicons of 200–220 nucleotides (nt) in length for three genes: 44 pairs of primers for *FOS*, 19 pairs of primers for *JUNB*, and 21 pairs of primers for *PLA2G2E* (Supplementary Table 2). For each probe, forward and reverse primers were synthesized by Sangon Biotech Company. Forward and reverse primer pairs were diluted to 100 µM in nuclease-free water. We carried out PCR reactions by mixing the following reagents: 5 µl of 10× rTaq buffer, 4 µl of 10× dNTP mix, 0.25 µl of rTaq Enzyme, 0.2 µl each of the 100 µM forward and reverse primers, 100ng of human genomic DNA, and nuclease-free water to a final volume of 50 µl. For each PCR reaction, 30 cycles were performed with Ta = 55 °C. After PCR, the contents of all the wells corresponding to a given probe were purified by agarose gel electrophoresis and gel recovery was carried out using HiPure Gel Pure DNA Mini Kit (Magen). Each of the amplicons was mixed together to generate an amplicon pool. 1ug of the 200 nt-220 nt amplicon pool was used for subsequent ethanol precipitation.

The *FOS*/*JUNB*/*PLA2G2E* PCR-based DNA FISH probes comprised 44/19/21 different 200-220 nt amplicon pools. For gene labeling, a volume corresponding to 1 µg from each 200-220 nt amplicon pool was precipitated by ethanol and then labeled with Alexa Fluor 647 using the ULYSIS Nucleic Acid Labeling Kit (Invitrogen, U21660), according to the supplier’s protocol. Unbound dyes were removed by Micro Bio-Spin Columns, P-30 Tris (Bio-Rad, 732-6223). The labeled probes were stored at −20 °C.

### Sample Preparation for ImmunoFISH

The sample was prepared for ImmunoFISH as previously described^58^, with minor modifications. For each hybridization, we used 2 ng/μl of the DNA probe in hybridization solution (namely, 100 ng of the FISH probe was precipitated and dissolved in 50 μl of the hybridization buffer). The hybridization buffer contained 50% formamide, 10% dextran sulfate, 2× SSC, and 1% Tween 20. Unlabeled competitor DNA (Cot-1 DNA and salmon sperm DNA) was added to the probes to inhibit nonspecific hybridization. The concentration of the competitor DNA was approximately 10 to 50 times that of the probe DNA. Therefore, we added 100 ng of the labeled probes, 10 μg of the unlabeled salmon sperm DNA, 1 μg of the unlabeled human Cot-1 DNA, and nuclease-free water to a final volume of 100 µl. Thereafter, 10 μl of 3 M sodium acetate and 275 μl of ice-cold 100% ethanol was added to the mixture. The labeled probe mix was precipitated at −80 °C for at least 1 hour. Efficient precipitation of the labeled probe mix was obtained by centrifuging at 14,800 rpm for 30 min. A second washing with 70% ethanol was then performed and the probe mix was centrifuged at 14,800 rpm for 5 min. The pellet was resuspended with hybridization mix consisting of 50% formamide, 10% dextran sulfate, 2× SSC, and 1% Tween 20. The hybridization mix was first denatured at 86 °C in a heat block for 2 min and immediately immersed on ice.

The cells were fixed with 4% paraformaldehyde in PBS for 10 min at room temperature. The cells were then washed twice with PBST (0.1% Tween 20 in PBS) and quenched with freshly prepared 20 mM glycine dissolved in PBS for 10 min. The cells were washed twice with PBST for 5 min each, permeabilized with 0.5% Triton X-100 and 0.5% saponin in PBS for 10 min, and then washed twice with PBST for 5 min each. The cells were incubated with 20% glycerol in PBS for at least 1 h at room temperature and then submerged into a liquid N2-filled container using fine forceps for approximately 6 sec. This freezing-thawing step was repeated twice to improve the penetration and accessibility of the probes to the target DNA sequences. The cells were washed twice with PBS for 2 min each. Thereafter, 0.1 M HCl was applied to the cells for 10 min at room temperature. The cells were washed twice with PBS for 2 min each and then washed twice with 2× SSC for 2 min each. The cells were incubated in 50% formamide and 2× SSC at 4 °C overnight. On the following day, the cells were pre-denatured in a water bath set at 73 °C for 10 min and then immediately placed on ice for 2 min. The cells were dehydrated through a series of 70%, 85%, and 100% ethanol baths for 1 min each and dried for 2 min. The pre-denatured probe hybridization mix was added to the dried cells. The cells were co-denatured with the probe on a heating block set at 86 °C for 3 min and then immediately placed on ice for 5 min. Finally, the samples were incubated in a hybridization oven set at 37 °C for at least 2 days. After hybridization, the samples were washed as follows: 3 times for 5 min each with 0.1% Tween 20 in 2× SSC at 37 °C on a shaking platform, 3 times for 5 min each with 0.3% Tween 20 in 0.1× SSC preheated to 63 °C under moderate shaking, 2 times for 5 min each with 0.1% Tween 20 in 4× SSC at 37 °C under moderate shaking, and 2 times for 5 min each with 2× SSC under moderate shaking.

The cells were incubated in PBST blocking buffer (2% BSA, 0.5% fish skin gelatin, and 0.02% Tween 20 in PBS) for 1 hour at room temperature. The primary antibody (mouse Pol II 8WG16, Abcam, ab817) was diluted 1:200 in PBST. The cells were incubated with the primary antibody in a dark humidified chamber for 3 hours. After incubation, the cells were washed 4 times with PBST. The secondary antibody (donkey anti-mouse, Cy3B) was diluted 1:20 in PBST. The samples were incubated with the secondary antibody in a dark humidified chamber for 45 min. After incubation, the cells were washed 4 times with PBST and then post-fixed in 4% paraformaldehyde in PBS for 10 min at room temperature. After post-fixation, the cells were washed 4 times with PBST and then mounted for imaging.

### Sample Preparation for Two-Color PALM/STORM Imaging of Nascent RNA and Pol II

The samples were prepared using Click-iT Plus Alexa Fluor 647 Picolyl Azide Toolkit (Molecular Probes, C10643), according to the supplier’s protocol. A 2 mM working solution of 5-ethynyl uridine (EU, Molecular Probes, E10345) was prepared from the 100 mM stock solution in pre-warmed complete medium. Half of the media within the 35-mm glass-bottom dish was replaced with fresh media containing 2 mM of EU. The cells were incubated under normal cell culture conditions for 10 min. The media were then aspirated, and the cells were fixed with 1 ml of 4% formaldehyde in PBS for 15 min at room temperature. Thereafter, the fixative was removed, and the cells were washed twice with 3% BSA in PBS, followed by permeabilization with 1 ml of 0.5% Triton X-100 in PBS for 20 min at room temperature. A 1× working solution of Click-iT reaction buffer additive was prepared by diluting the 10× solution 1:10 in deionized water. The CuSO4 and copper protectant pre-mix was prepared by mixing 5 μl of CuSO4 with 5 μl of copper protectant. The Click-iT Plus reaction cocktail was prepared by mixing 435 μl of 1× Click-iT Plus reaction buffer, 5 μl of 500 μM Alexa Fluor 647 PCA solution, 10 μl of CuSO4-copper protectant pre-mix, and 50 μl of 1× Click-iT reaction buffer additive. The cells were washed twice with 3% BSA in PBS. 500 μl of Click-iT Plus reaction cocktail was added to the cells. The dish was wrapped with aluminum foil and incubated for 30 min at room temperature. Thereafter, the sample was washed once with 3% BSA in PBS and immediately mounted for imaging.

### Spinning Disk Confocal Microscopy

Spinning disk confocal microscopy was conducted on an UltraVIEW VoX spinning disk microscope (PerkinElmer) with an EMCCD (Hamamatsu C9100-13). Single z-slice images were acquired at an exposure time of 100 ms for both 514 nm and 440 nm laser lines. Image analysis was performed using Volocity software.

### Sample Preparation for Two-Color PALM/STORM Imaging of Nuclear Actin and Pol II

Cells cultured on a 35-mm glass-bottom dish were briefly washed twice with PBS and then permeabilized with 1ml of hypotonic buffer (10 mM Tris-HCl pH 7.3, 10 mM KCl, and 1.5 mM MgCl2) for 3 min. Thereafter, the cells were washed with 500 μl of 0.5% PBST (0.5% Triton X-100 in PBS) for 45 sec, followed by a second washing with PBST for 30 sec and a third washing with PBS. The cells were fixed with 1 ml of 4% paraformaldehyde (EM grade, Electron Microscropy Sciences) in cytoskeleton buffer (10 mM MES pH 6.1, 150 mM NaCl, 5 mM EGTA, 5 mM glucose, and 5 mM MgCl2) for 10 min. The cells were then washed with PBS three times for 10 min each. To label each dish of sample, the staining solution was prepared by diluting 30 μl of the Alexa Fluor 647 Phalloidin (Molecular Probes, A22287) stock solution (∼6.6 μM in methanol) into 400 μl of PBS (a final concentration of ∼0.5 μM). The dishes were wrapped with aluminum foil and incubated at 4 °C overnight on a shaker. Thereafter, the sample was briefly washed once with PBS and immediately mounted for imaging.

### Epifluorescence, tcPALM, and Two-Color PALM/STORM Imaging

Epifluorescence, tcPALM, and two-color PALM/STORM imaging were conducted using an Olympus inverted microscope equipped with a 100× oil-immersion objective (Olympus, NA 1.49) and oblique incidence excitation. A 488 nm laser (MPB Communications, 500 mW) was used to excite the fluorophore Dendra2 for conventional epifluorescence imaging. A 405 nm laser (Coherent, Cube 405-100C) was used for the photoconversion of Dendra2 from the green-emission state to the red-emission state or for the activation of Alexa Fluor 647 from the dark state to the fluorescent state. A 561 nm laser (MPB Communications, 2 W) was used to illuminate Dendra2 at the red-emission state. A 647 nm laser (MPB Communications, 1.5 W) was used to excite Alexa Fluor 647. The laser lines were combined with a custom dichroic (Chroma, zt405/488/561/647/752RPC). The emission was filtered with a custom dichroic (Chroma, zet405/488/561/647-656/752M). The emission was separated with a QuadView (Photometrics, QV2) using the dichroics T560lpxr, T650lpxr, and 750dcxxr (Chroma) and the emission filters ET525/50m, ET605/70m, ET700/75m, and HQ795/50m (Chroma), and imaged with an EMCCD (Andor-897). The z-position was maintained during acquisition by detecting a separate IR beam reflected from the coverslip using a quadrant photodiode, which reads the position of the coverslip and feeds back the position to a piezo-controlled objective.

For epifluorescence imaging, images were acquired at an exposure time of 200 ms and an EM gain of 30 for 5 consecutive frames (the 5 frames were averaged for image analysis). The laser power density on the sample was 0.2 W/cm^2^ (488 nm).

For tcPALM imaging, images were acquired at exposure time of 50 ms and an EM gain of 90 for 10,000 frames under continuous illumination for both 405 nm and 561 nm lasers. The laser power densities on the sample were 0.15 W/cm^2^ (405 nm) and 2.0 kW/cm^2^ (561 nm).

For two-color PALM/STORM imaging, images of Alexa Fluor 647 were acquired at an exposure time of 20 ms and an EM gain of 30 for ∼ 20,000 frames under continuous illumination for a single 647 nm laser, then images of Dendra2 were acquired at an exposure time of 50 ms and an EM gain of 90 for ∼ 10,000 frames under continuous illumination for both 405 nm and 561 nm lasers. Fiducial beads for drift correction and chromatic correction were simultaneously tracked during image acquisition. The laser power densities on the sample were 1.5 kW/cm^2^ (647 nm) and 2.0 kW/cm^2^ (561 nm). The 405 nm laser intensity on the sample was ramped from 0.1 W/cm^2^ to 30 W/cm^2^ to maintain an almost constant rate of photoswitching. The cells were imaged in STORM imaging buffer. The STORM imaging buffer was prepared by mixing 2 ml of 20 mM NaCl, 100 mM Tris pH 8.0, 10% (w/v) glucose with 20 μl Glucose Oxidase Oxygen Scavenger solution and 20 μl β-mercaptoethanol (Sigma-Aldrich 63689-25ML-F). The Glucose Oxidase Oxygen Scavenger solution contained 60 mg/ml glucose oxidase (Sigma-Aldrich G6125-10KU), 6 mg/mL catalase from bovine liver (Sigma-Aldrich C100-50MG), and 40% glycerol in PBS.

Image analysis was conducted using Insight3 software (provided by Bo Huang from University of California, San Francisco) for fluorophore localization and custom-written Matlab scripts for drift correction, chromatic correction, and image reconstruction.

### Structured Illumination Microscopy (SIM) Imaging for ImmunoFISH

SIM imaging was performed under an N-SIM microscope (Nikon) equipped with a 100× oil-immersion objective (Nikon, NA 1.49) and an EMCCD (Andor, iXon3 DU-897E). A z-stack of 7 layers with a step size of 0.12 μm was acquired at an exposure time of 20-200 ms, an EM gain of 300, and a conversion gain of 2.4× for 640 nm and 561 nm laser lines. The exposure time and laser intensities were adjusted to achieve a good image quality. SIM images were reconstructed by the 3D Reconstruct Stack program in the N-SIM module. The reconstructed z-stack image was projected into a 2D image by maximum intensity projection. The cells were imaged in STORM imaging buffer without β-mercaptoethanol.

### Cluster Analysis

Clusters were identified using open-source SR-Tesseler software^28^. Clusters smaller than 40 nm in diameter (1256.637 nm^2^ in area) were excluded from our analysis. Pair auto-correlation analysis and tcPALM analysis were performed as previously described^17, 27^ using custom-written Matlab scripts. For tcPALM analysis, localizations separated by no more than 30 dark frames were considered part of the same burst; bursts with fewer than 10 localizations were excluded from our analysis as such a burst might arise from the blinking of a single Dendra2 molecule.

tcPALM is a method to capture and quantify transient clustering dynamics in living cells. For a Pol II cluster in living cells, single-molecule signals appeared clustered in the detection profile and exhibited large steps in the cumulative count (Supplementary Fig. 6, left). This large step represented an abrupt increase in local Pol II concentration, indicating the assembly of a Pol II cluster. This assembly event was termed a burst. The number of localizations in a burst was termed burst size and the duration of a burst was termed burst lifetime (Supplementary Fig. 6, left). On the contrary, for a Pol II cluster in fixed cells, single-molecule signals concentrated at the start of the acquisition and showed a monotonic slope gradually followed by a plateau (Supplementary Fig. 6, right). This curve represented the gradual photobleaching of immobile Dendra2-Pol II in fixed cells.

### Colocalization Analysis

Colocalization of two channels in two-color PALM/STORM imaging was analyzed by calculating the percentage of cluster area overlap (area overlap ratio) and analyzing the pair cross-correlation. For the calculation of the area overlap ratio, the clusters in the two channels were identified by SR-Tesseler^28^. The two cluster images were exported from SR-Tesseler and then opened in ImageJ software (National Institutes of Health, USA). The colocalization analysis was conducted on a cluster-to-cluster basis for the area of nucleoplasm (the regions of nucleoli and nuclear membrane were excluded from the analysis). The total areas of Pol II clusters, actin clusters, and overlap between these two clusters were calculated by the particle analyzer in ImageJ. The area overlap ratio was calculated as follows:

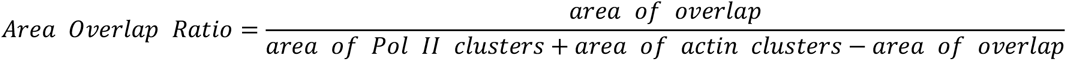

Pair cross-correlation analysis was performed as previously described^27^ using custom-written Matlab scripts.

### Statistics and Reproducibility

Statistical analyses were conducted using OriginPro 8 software. Statistical significance was determined by one-way ANOVA with Tukey–Kramer test, two-tailed *t*-test, and pair-sample two-tailed *t*-test as indicated in the corresponding figure legends.

All experiments presented in the manuscript were repeated at least in two independent experiments/biological replicates.

### Data Availability

RNA-Seq data have been deposited at Gene Expression Omnibus (GEO) under the accession code GSE82320.

(http://www.ncbi.nlm.nih.gov/geo/query/acc.cgi?token=azqrckqgdfqbpgh&acc=GSE82320)

