## Supplementary Information for "Nuclear actin regulates inducible transcription by enhancing RNA polymerase II clustering"

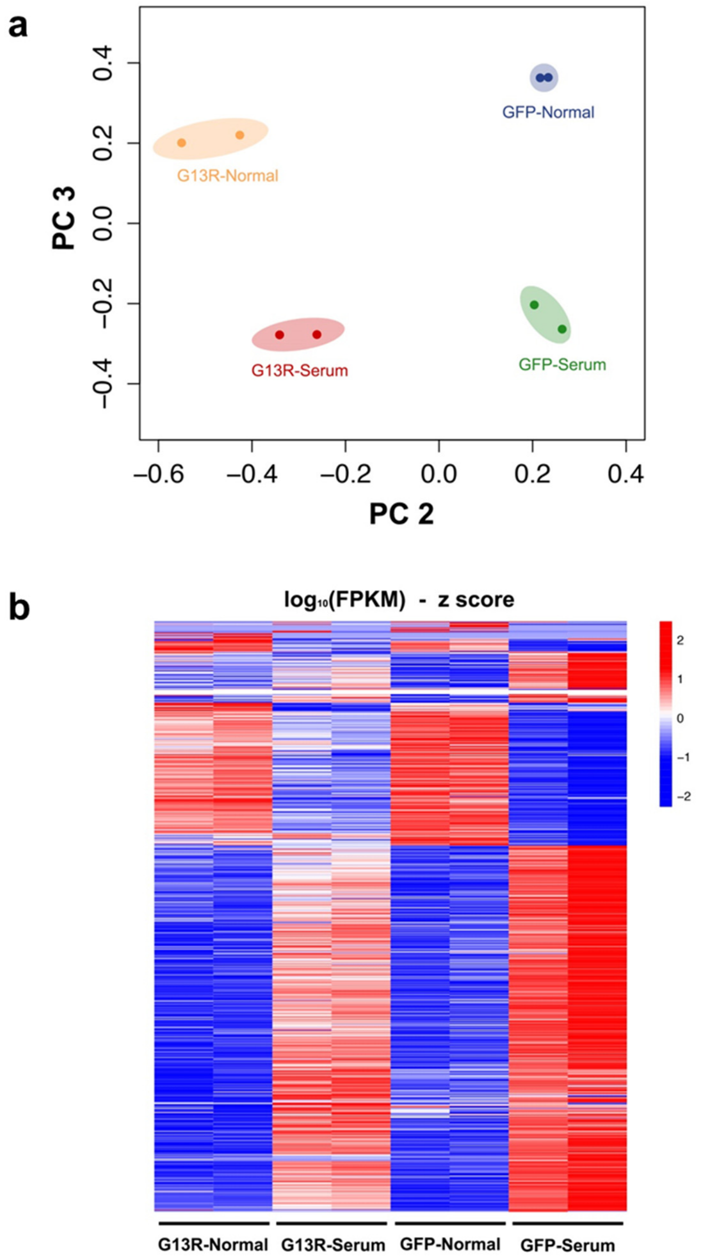


**Supplementary Fig. 1 | Principal component analysis and differential gene expression analysis show great consistency between replicates.**

**a**, Principal component analysis of RNA-seq data. **b**, Heatmap showing FPKM (z score) of differentially expressed genes in cells overexpressing G13R and cells overexpressing GFP under normal-growth and serum-stimulation conditions.

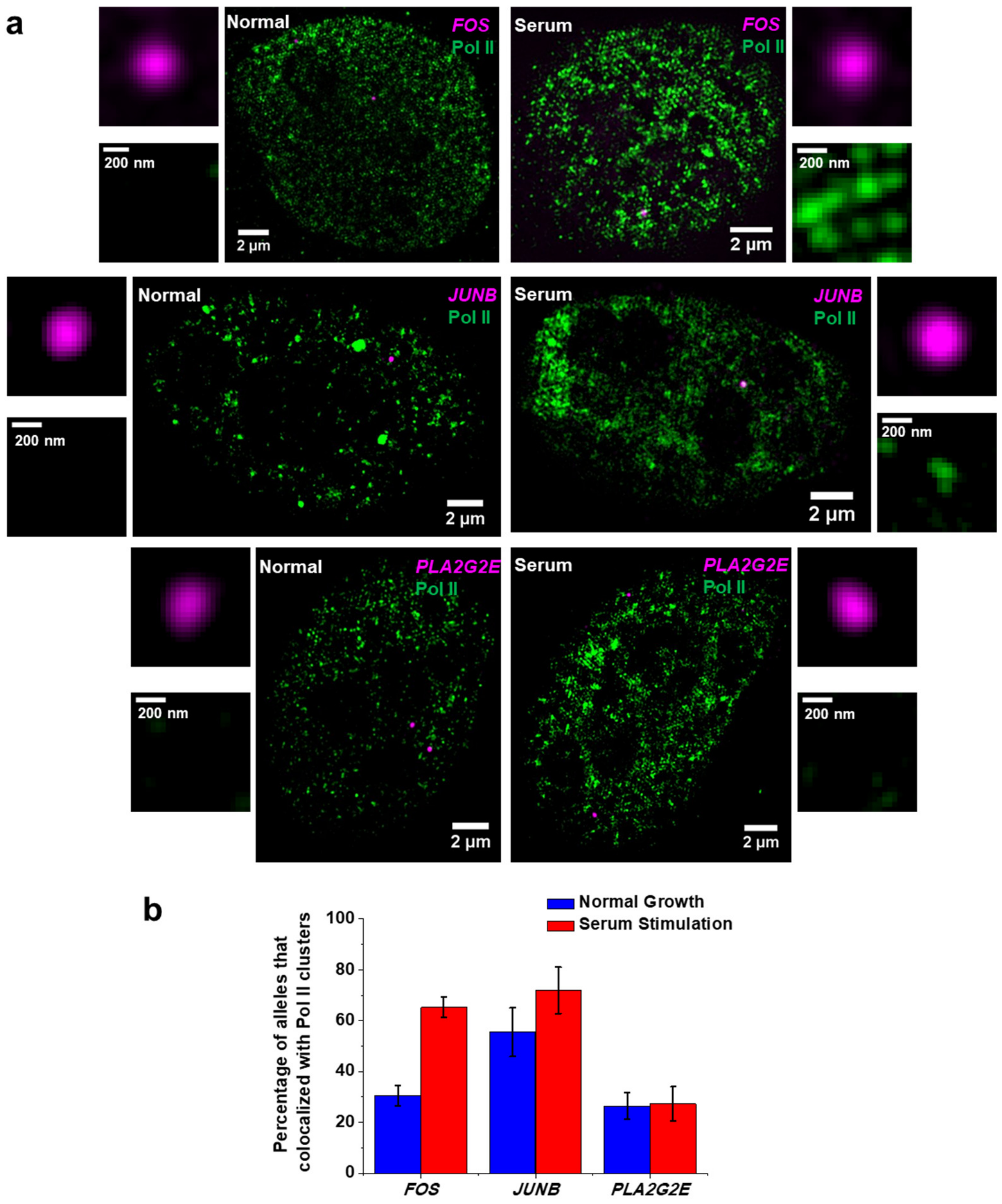


**Supplementary Fig. 2 | Specific serum-response genes are localized within Pol II clusters upon serum stimulation.**

**a**, Representative Structured Illumination Microscopy (SIM) ImmunoFISH images showing Pol II (green) and gene loci (magenta) of *FOS* (top, n = 60 cells under normal-growth condition, n = 65 cells under serum-stimulation condition), *JUNB* (middle, n = 20 cells under normal-growth condition, n = 20 cells under serum-stimulation condition), and *PLA2G2E* (bottom, n = 40 cells under normal-growth condition, n = 26 cells under serum-stimulation condition). Zoom-in images of gene loci and Pol II are shown near the corresponding images of whole nuclei. **b**, Percentage of alleles that colocalized with Pol II clusters in cells under normal-growth and serum-stimulation conditions. (*FOS*: n = 60 cells under normal-growth condition, n = 65 cells under serum-stimulation condition. *JUNB*: n = 20 cells under normal-growth condition, n = 20 cells under serum-stimulation condition. *PLA2G2E*: n = 40 cells under normal-growth condition, n = 26 cells under serum-stimulation condition). Data are shown as mean ± SD (bootstrapping).

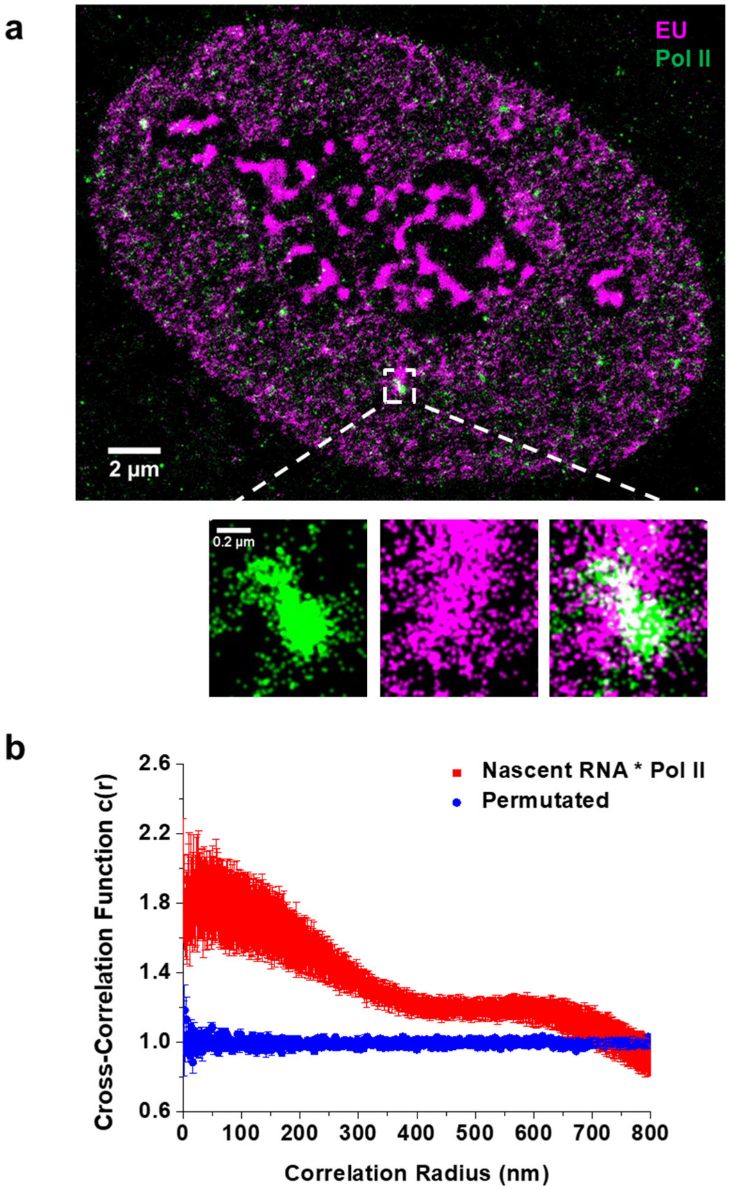


**Supplementary Fig. 3 | Pol II clusters are active transcription sites.**

**a**, Representative two-color PALM/STORM images showing nascent RNA (magenta) and Pol II (green) (n = 7 cells). Zoom-in images of the selected region are shown below the full-size image. **b**, Quantification of colocalization by pair cross-correlation analysis. Data are shown as mean ± SEM. (n = 7 cells for original images, n = 7 for pixel-permutated images)

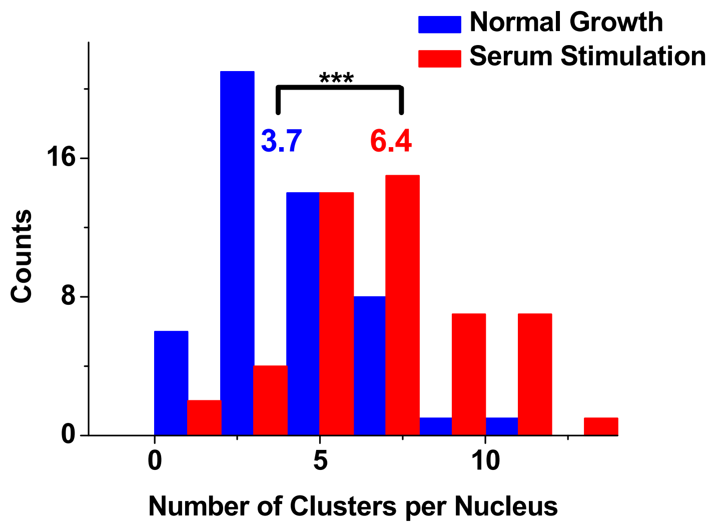


**Supplementary Fig. 4 | Increased number of Pol II clusters upon serum stimulation revealed by epifluorescence microscopy.**

Distribution of the number of clusters per nucleus under normal-growth (n = 50 cells) and serum-stimulation (n = 50 cells) conditions under epifluorescence microcopy. Statistical significance was determined by two-tailed *t*-test. *$p < 0.05$, **$p < 0.01$, and ***$p < 0.001$.

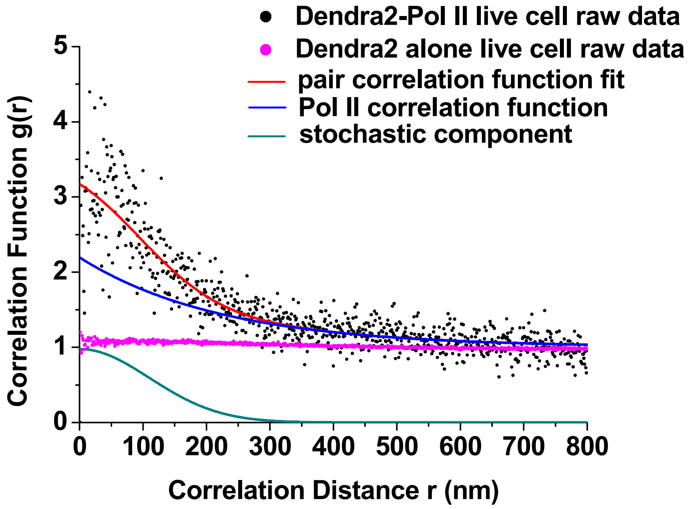


**Supplementary Fig. 5 | Pol II clustering revealed by pair correlation analysis.**

Representative pair correlation analysis of the U2OS cell line stably expressing Dendra2-Pol II (black dots) and normal U2OS cells transiently expressing Dendra2 (pink dots). The correlation curve of Dendra2-Pol II was fit to a fluctuation model (red line) that separated Pol II correlation function (blue line) from the stochastic component due to blinking of Dendra2 (green line).

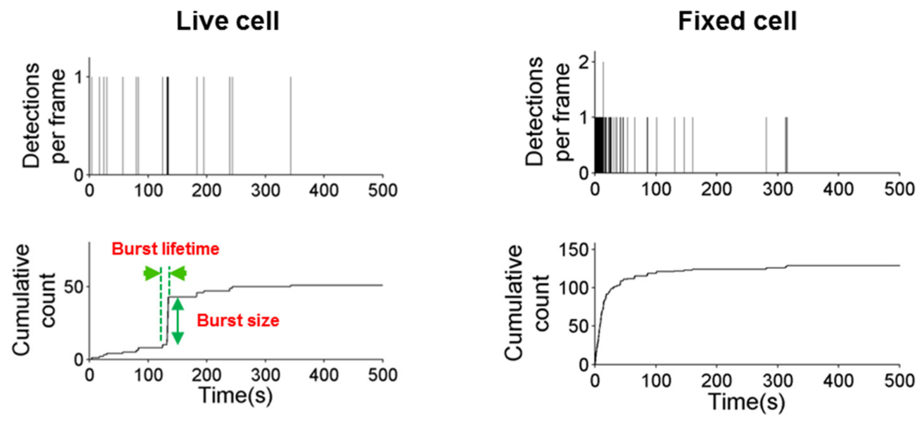


**Supplementary Fig. 6 | Dynamic clustering of Pol II revealed by tcPALM.**

Representative tcPALM profiles of a Pol II cluster in a live cell (left) and a Pol II cluster in a fixed cell (right).

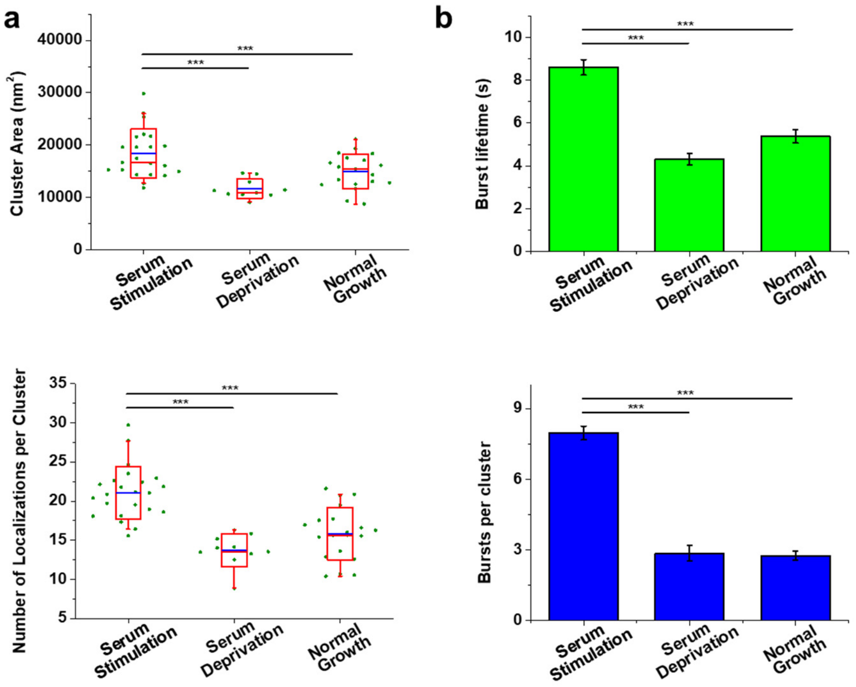


**Supplementary Fig. 7 | Enhanced Pol II clustering upon serum stimulation.**

**a**, Cluster area and the number of localizations per cluster under serum-stimulation (n = 23 cells), serum-deprivation (n = 10 cells), and normal-growth conditions (n = 20 cells). **b**, Burst lifetime and the number of bursts per cluster under serum-stimulation (n = 829 bursts of 104 clusters from 23 cells), serum-deprivation (n = 174 bursts of 61 clusters from 10 cells), and normal-growth conditions (n = 306 bursts of 111 clusters from 20 cells). For boxplots, bar charts, and statistics see the legend of Fig. 3.

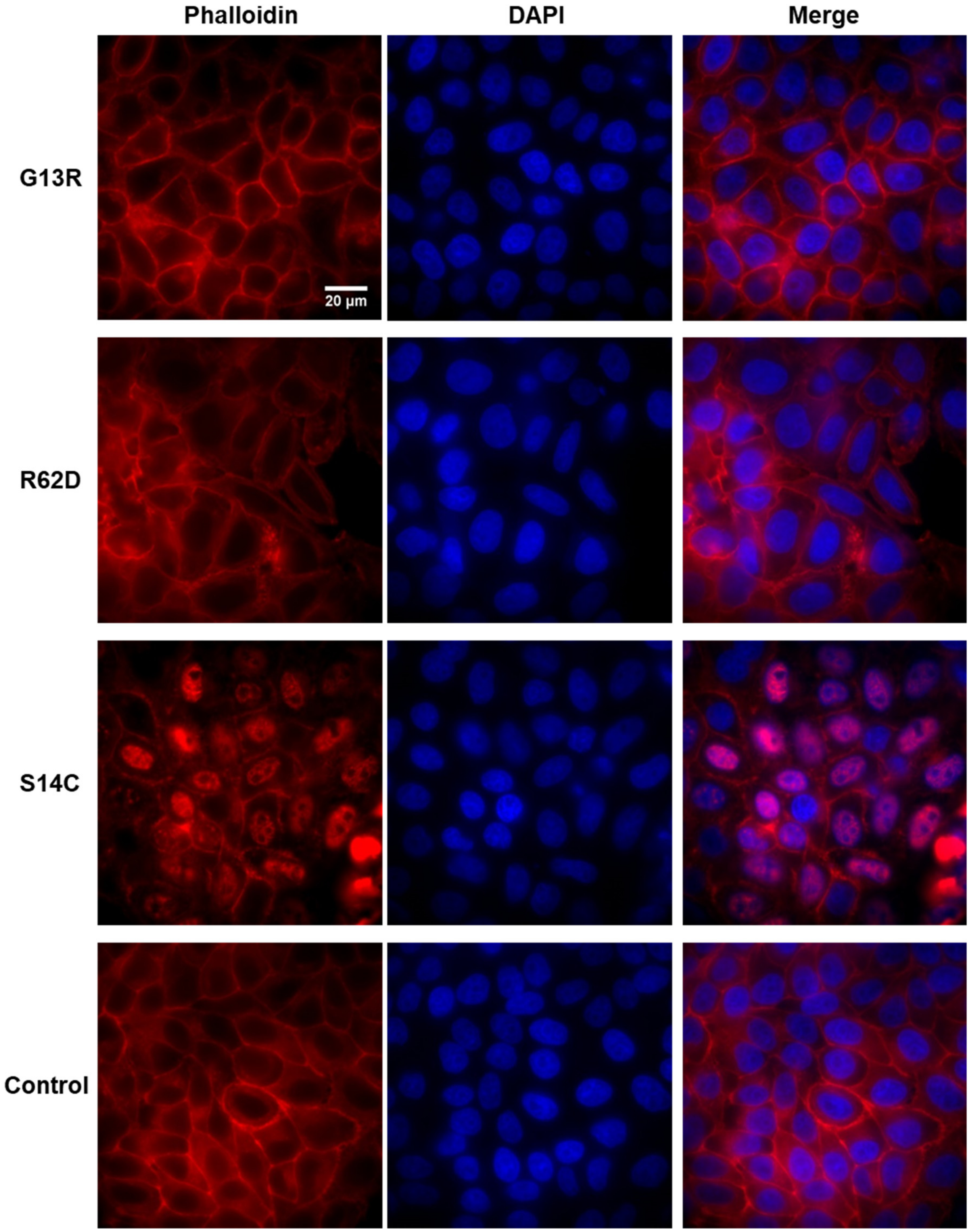


**Supplementary Fig. 8 | Characterization of actin mutants.**

Representative epifluorescence images showing phalloidin-labeled F-actin (left), DAPI-labeled nuclei (middle), and the merge (right) in cells overexpressing actin mutants and control cells.

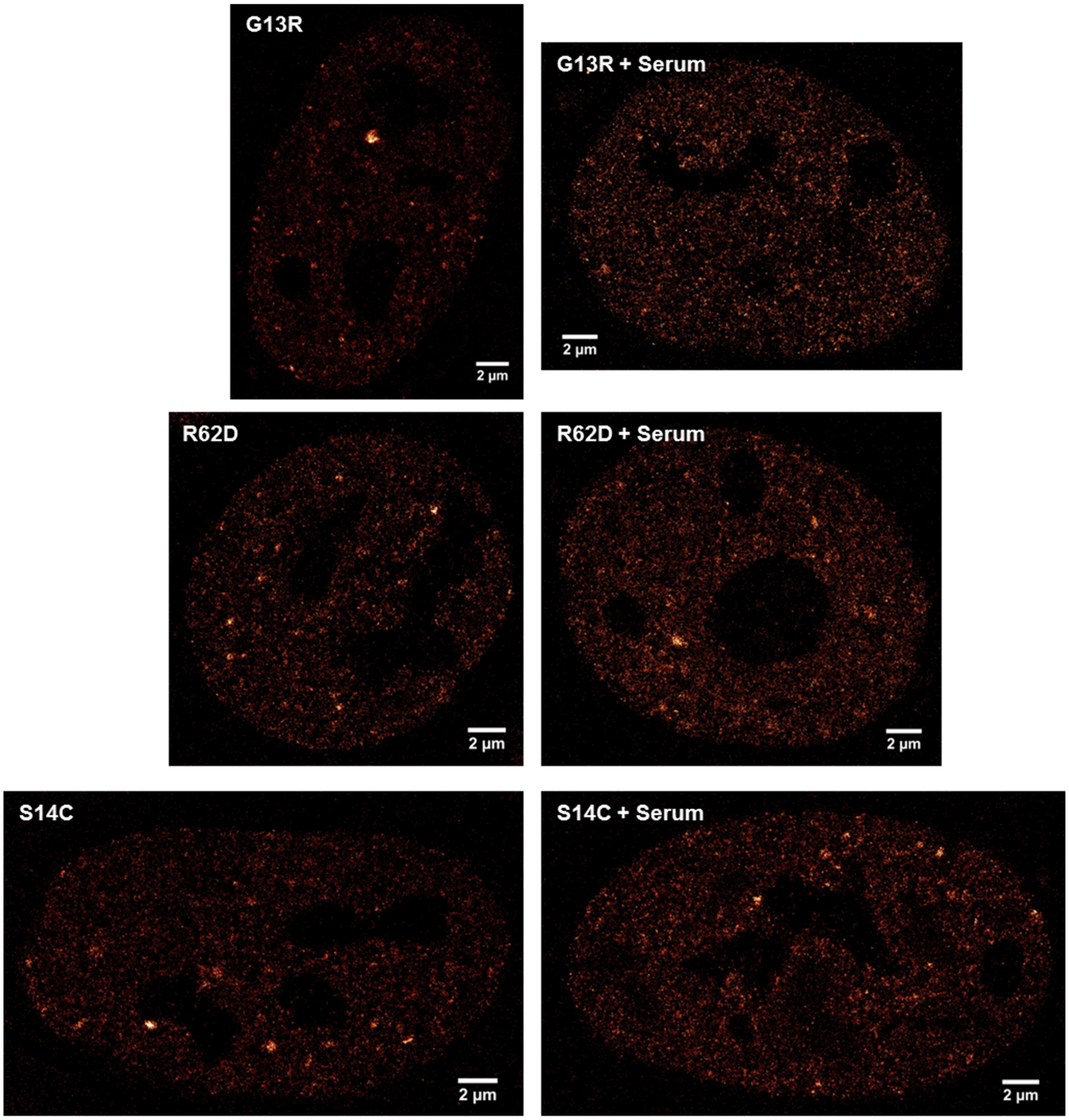


**Supplementary Fig. 9 | PALM images of Pol II clusters in cells overexpressing actin mutants.**
Representative PALM images of Pol II clusters in cells overexpressing actin mutants G13R, R62D, and S14C (from top to bottom) under normal-growth (left) and serum-stimulation (right) conditions.

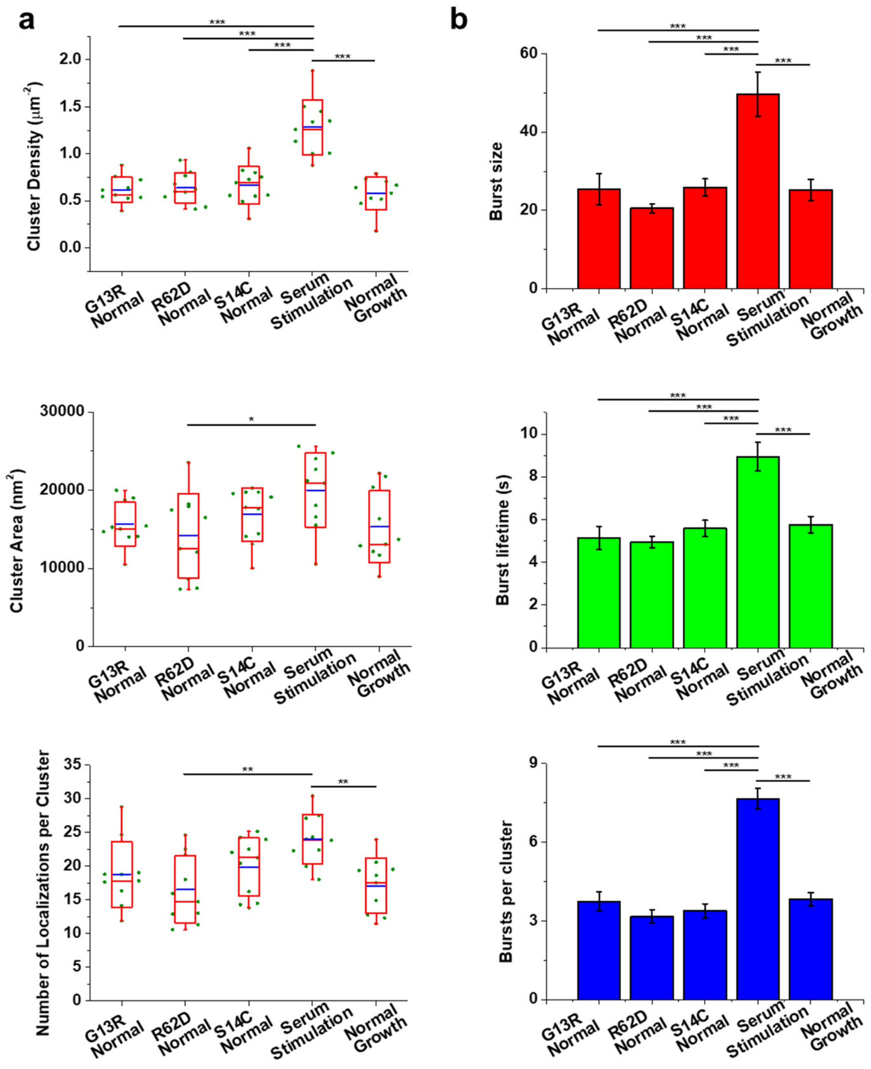


**Supplementary Fig. 10 | Overexpression of actin mutants does not affect Pol II clustering under normal-growth condition.**

**a**, Cluster density, cluster area, and the number of localizations per cluster in cells overexpressing different actin mutants under normal-growth condition (n = 10, 10, and 11 cells, respectively, from left to right), cells under serum-stimulation condition (n = 10 cells), and cells under normal-growth condition (n = 10 cells). **b**, Burst size, burst lifetime, and the number of bursts per cluster in cells overexpressing different actin mutants under normal-growth condition (n = 232 bursts of 62 clusters from 10 cells, 216 bursts of 68 clusters from 10 cells, and 240 bursts of 71 clusters from 11 cells, respectively, from left to right), cells under serum-stimulation condition (n = 406 bursts of 53 clusters from 10 cells), and cells under normal-growth condition (n = 287 bursts of 75 clusters from 10 cells). For boxplots, bar charts, and statistics see the legend of Fig. 3.

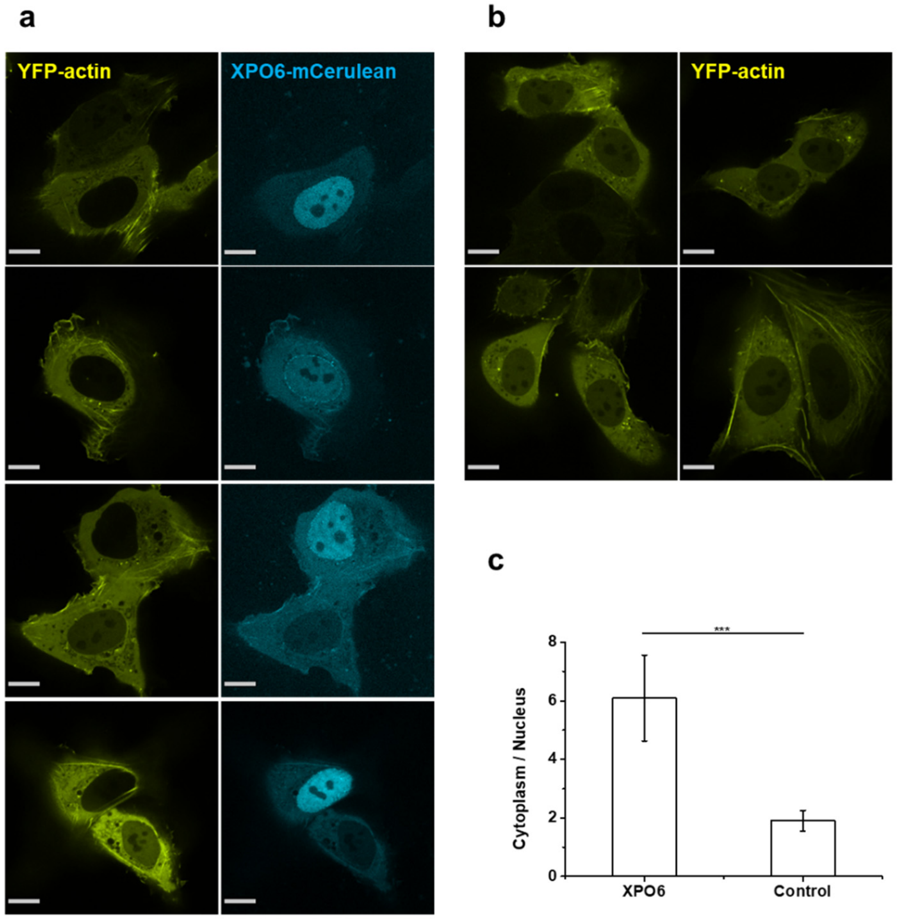


**Supplementary Fig. 11 | Overexpression of XPO6 significantly reduces nuclear actin.**

**a**, Representative spinning-disk confocal images of cells cotransfected with YFP-actin (yellow) and XPO6-mCerulean (cyan). **b**, Representative spinning-disk confocal images of cells transfected with YFP-actin (yellow). **c**, The ratio of fluorescence in the cytoplasm to that in the nucleus in cells expressing both YFP-actin and XPO6-mCerulean ("XPO6", n = 36 cells) and control cells expressing only YFP-actin ("Control", n = 44 cells). Data are shown as mean ± SD. Statistical significance was determined by two-tailed *t*-test. *$p< 0.05$, **$p< 0.01$, and ***$p< 0.001$. Scale bar is 10 μm in a and b.

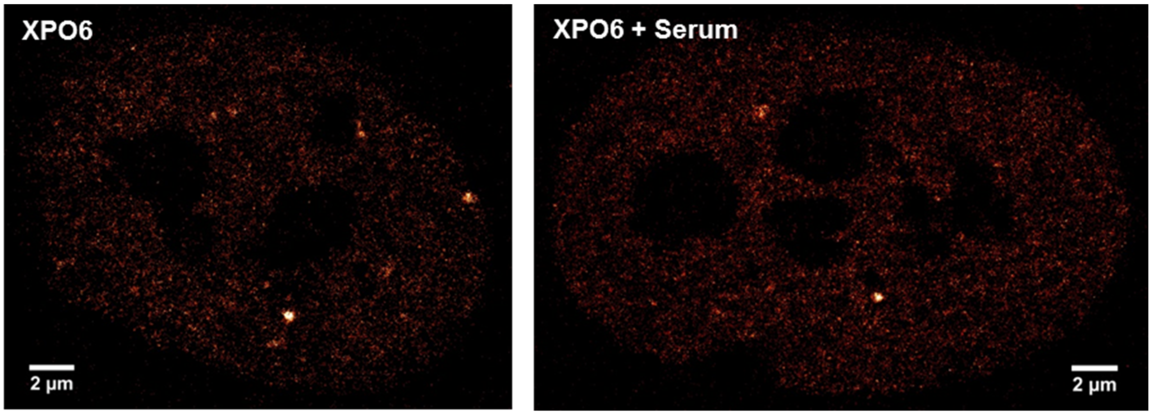


**Supplementary Fig. 12 | PALM images of Pol II clusters in cells overexpressing XPO6.**
Representative PALM images of Pol II clusters in cells overexpressing XPO6 under normal-growth (left) and serum-stimulation (right) conditions.

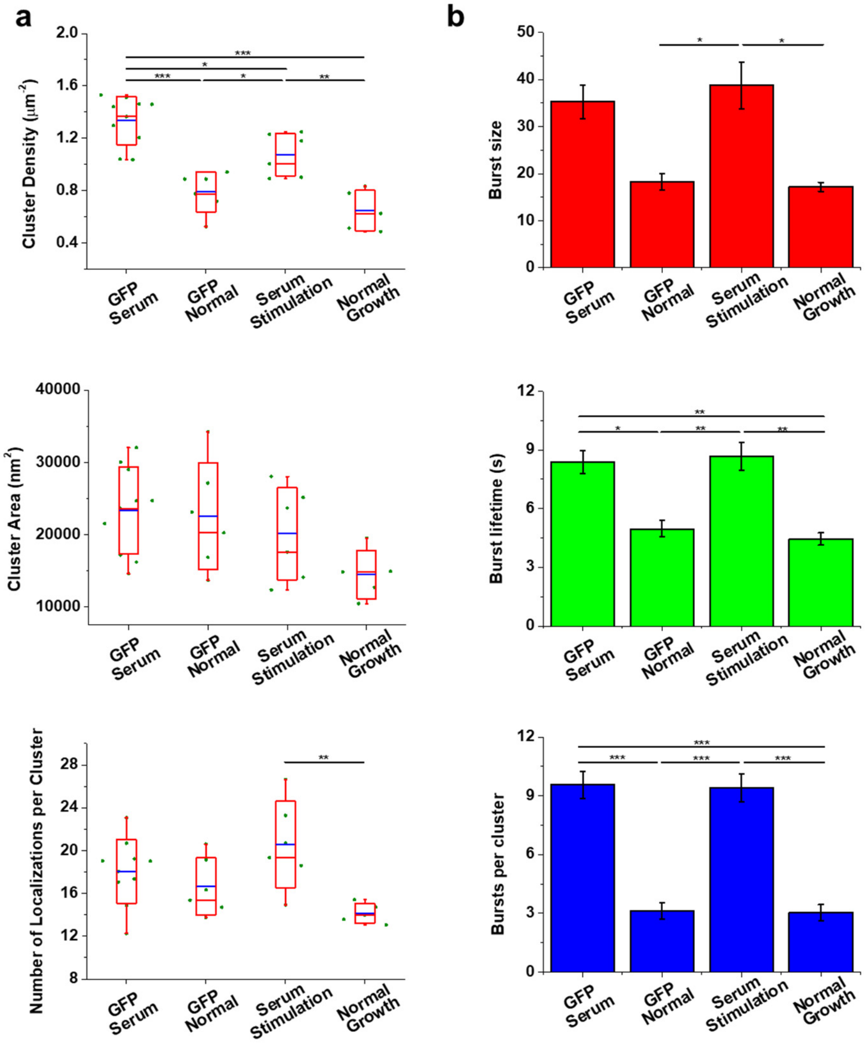


**Supplementary Fig. 13 | Transfection procedure does not interfere with the enhanced-level Pol II clustering upon serum stimulation.**

**a**, Cluster density, cluster area, and the number of localizations per cluster in cells overexpressing GFP under serum-stimulation (n = 10 cells) and normal-growth (n = 6 cells) conditions, cells under serum-stimulation condition (n = 6 cells), and cells under normal-growth condition (n = 5 cells). **b**, Burst size, burst lifetime, and the number of bursts per cluster in cells overexpressing GFP under serum-stimulation (n = 325 bursts of 34 clusters from 10 cells) and normal-growth (n = 94 bursts of 30 clusters from 6 cells) conditions, cells under serum-stimulation condition (n = 235 bursts of 25 clusters from 6 cells), and cells under normal-growth condition (n = 76 bursts of 25 clusters from 5 cells). For boxplots, bar charts, and statistics see the legend of Fig. 3.

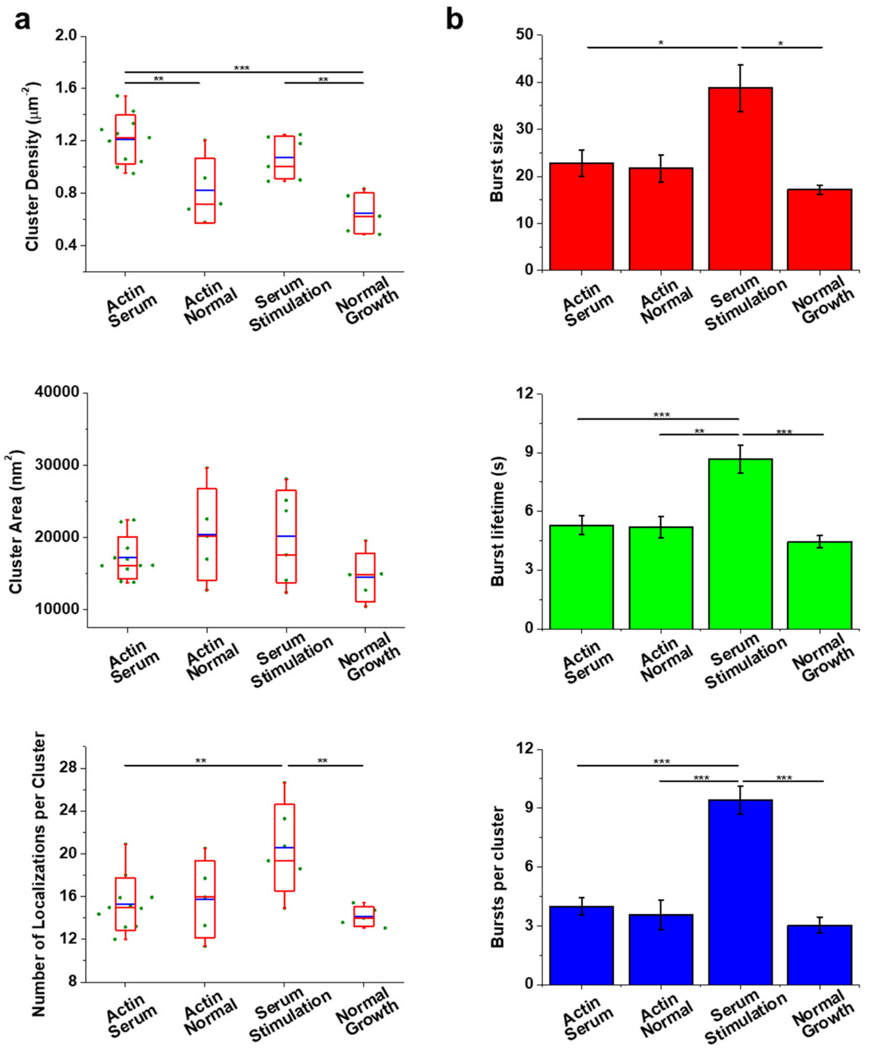


**Supplementary Fig. 14 | Overexpression of wild-type actin in the nucleus does not affect the spatial organization but interferes with the temporal dynamics, of Pol II clusters.**

**a**, Cluster density, cluster area, and the number of localizations per cluster in cells overexpressing NLS-wild-type actin under serum-stimulation (n = 11 cells) and normal-growth (n = 5 cells) conditions, cells under serum-stimulation condition (n = 6 cells), and cells under normal-growth condition (n = 5 cells). **b**, Burst size, burst lifetime, and the number of bursts per cluster in cells overexpressing NLS-wild-type actin under serum-stimulation (n = 148 bursts of 37 clusters from 11 cells) and normal-growth (n = 89 bursts of 25 clusters from 5 cells) conditions, cells under serum-stimulation condition (n = 235 bursts of 25 clusters from 6 cells), and cells under normal-growth condition (n = 76 bursts of 25 clusters from 5 cells). The data for cells under serum-stimulation and normal-growth conditions are the same as those in Supplementary Fig. 13. For boxplots, bar charts, and statistics see the legend of Fig. 3.

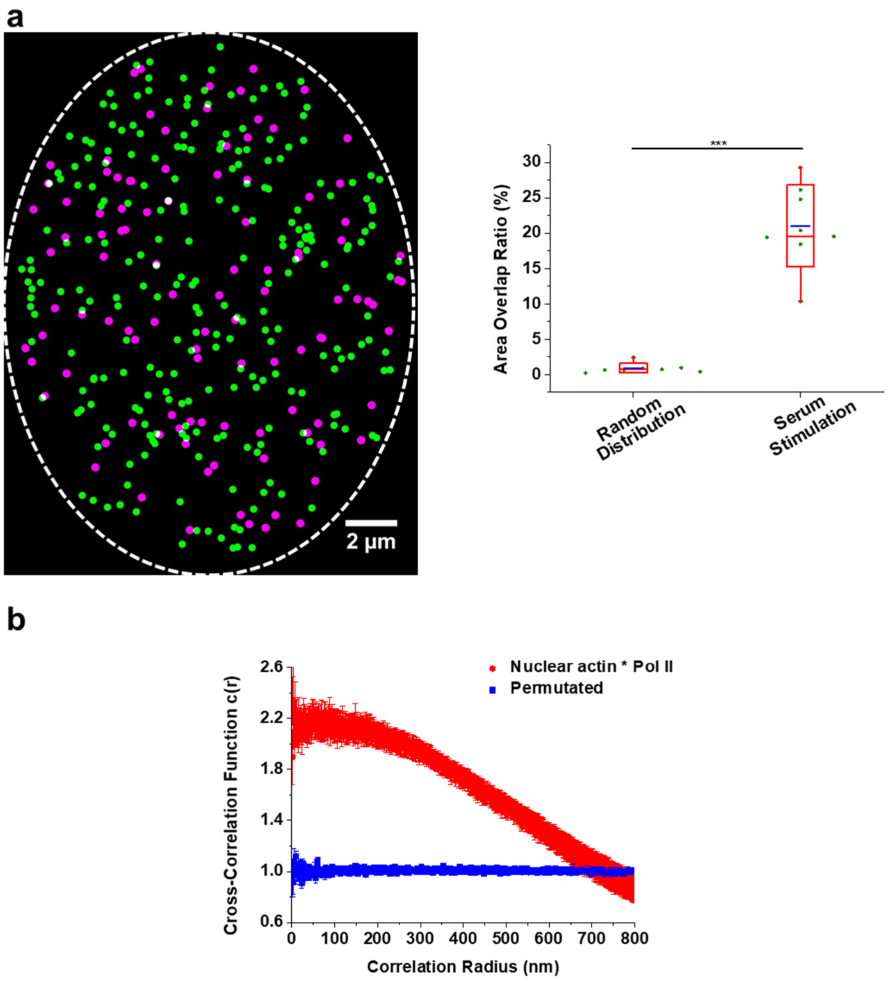


**Supplementary Fig. 15 | Colocalization between nuclear actin filaments and Pol II upon serum stimulation is higher than the random level.**

**a**, Simulation of randomly distributed clusters in an oval with the same cluster density and cluster area as those in cells upon serum stimulation. Left: a representative cluster image simulating nuclear actin clusters (magenta) and Pol II clusters (green). White dash lines delineate the cell nucleus. Right: quantification of colocalization by calculating the area overlap ratio (n = 8 cells). The data of "Serum Stimulation" is the same as that in Fig. 5B, left. Statistical significance was determined by pair-sample two-tailed *t*-test. **p*< 0.05, ***p*< 0.01, and ****p*< 0.001. **b**, Quantification of colocalization between nuclear actin filaments and Pol II by pair cross-correlation analysis. Data are shown as mean ± SEM. (n = 7 cells upon serum stimulation for original images, n = 7 for pixel-permutated images)

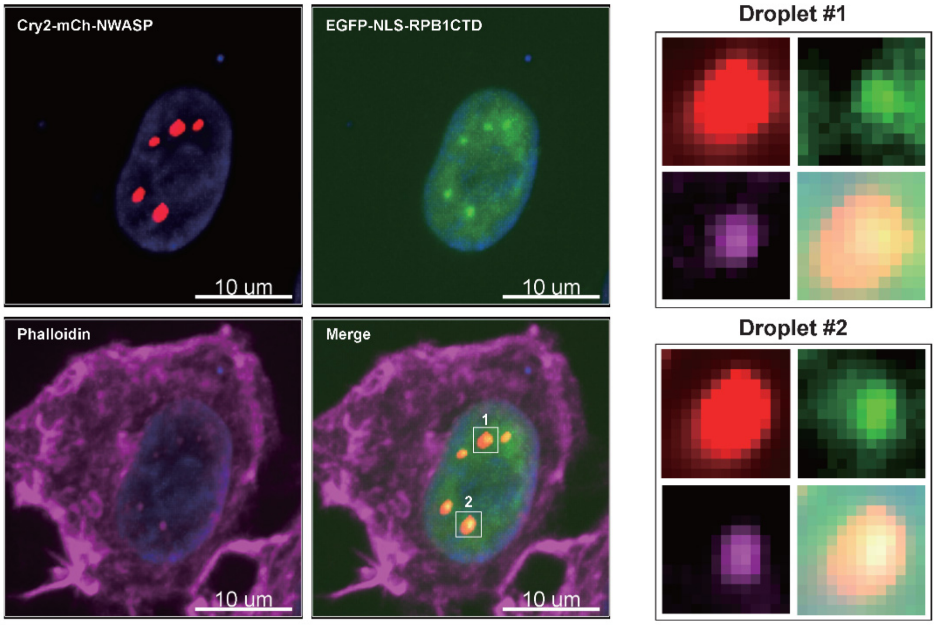


**Supplementary Fig. 16 | N-WASP phase-separates with the C-terminal domain (CTD) of RPB1 and nuclear actin.**

Left: representative images show that N-WASP phase-separated droplets incorporate the CTD of RPB1 and nuclear actin. Right: Zoom-in images of two droplets. Red, Cry2-mCherry-N-WASP. Blue, DAPI. Green, EGFP-NLS-RPB1 CTD. Purple, Phalloidin.

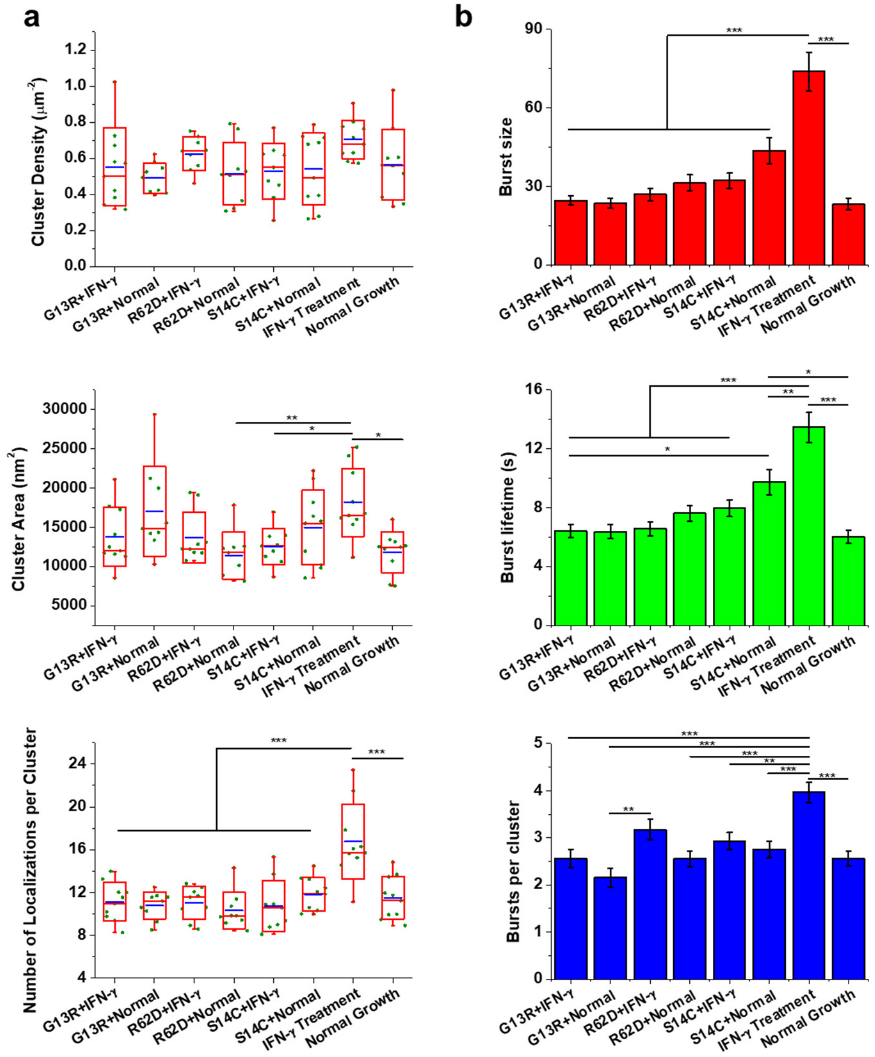


**Supplementary Fig. 17 | Overexpression of actin mutants abolishes enhanced-level Pol II clustering upon IFN-γ treatment.**

**a**, Cluster density, cluster area, and the number of localizations per cluster in cells overexpressing G13R under IFN-γ treatment and normal-growth conditions (n = 10 and 9 cells, respectively), cells overexpressing R62D under IFN-γ treatment and normal-growth conditions (n = 9 and 9 cells, respectively), cells overexpressing S14C under IFN-γ treatment and normal-growth conditions (n = 9 and 10 cells, respectively), cells under serum-stimulation condition (n = 10 cells), and cells under normal-growth condition (n = 10 cells). **b**, Burst size, burst lifetime, and the number of bursts per cluster in cells overexpressing G13R under IFN-γ treatment and normal-growth conditions (n = 128 bursts of 50 clusters from 10 cells and n = 97 bursts of 45 clusters from 9 cells, respectively), cells overexpressing R62D under IFN-γ treatment and normal-growth conditions (n = 146 bursts of 46 clusters from 9 cells and n = 115 bursts of 45 clusters from 9 cells, respectively), cells overexpressing S14C under IFN-γ treatment and normal-growth conditions (n = 132 bursts of 45 clusters from 9 cells and n = 138 bursts of 50 clusters from 10 cells, respectively), cells under serum-stimulation condition (n = 218 bursts of 55 clusters from 10 cells), and cells under normal-growth condition (n = 128 bursts of 50 clusters from 10 cells). For boxplots, bar charts, and statistics see the legend of Fig. 3.

**Supplementary Tables** (provided as separate Excel files)

**Supplementary Table 1** Differentially expressed genes and the corresponding functional annotations.

**Supplementary Table 2** Primers for the PCR-based synthesis of FISH probes.
